# Two transcriptionally distinct pathways drive female development in a reptile with both genetic and temperature dependent sex determination

**DOI:** 10.1101/2021.02.03.429474

**Authors:** Sarah L. Whiteley, Clare E. Holleley, Susan Wagner, James Blackburn, Ira W. Deveson, Jennifer A. Marshall Graves, Arthur Georges

**Affiliations:** Institute for Applied Ecology, University of Canberra, Australia; Australian National Wildlife Collection CSIRO National Research Collections Australia, Canberra, Australia; Garvan Institute of Medical Research, Sydney, Australia; St. Vincent’s Clinical School, UNSW, Sydney, Australia; Latrobe University, Melbourne, Australia

**Keywords:** Sex reversal, gonad differentiation, thermosensitivity, calcium signalling, oxidative stress

## Abstract

How temperature determines sex remains unknown. A recent hypothesis proposes that conserved cellular mechanisms (calcium and redox; ‘CaRe’ status) sense temperature and identify genes and regulatory pathways likely to be involved in driving sexual development. We take advantage of the unique sex determining system of the model organism, *Pogona vitticeps*, to assess predictions of this hypothesis. *P. vitticeps* has ZZ male: ZW female sex chromosomes whose influence can be overridden in genetic males by high temperatures, causing male-to-female sex reversal. We compare a developmental transcriptome series of ZWf females and temperature sex reversed ZZf females. We demonstrate that early developmental cascades differ dramatically between genetically driven and thermally driven females, later converging to produce a common outcome (ovaries). We show that genes proposed as regulators of thermosensitive sex determination play a role in temperature sex reversal. Our study greatly advances the search for the mechanisms by which temperature determines sex.

**Author Summary:** In many reptiles and fish, environment can determine, or influence, the sex of developing embryos. How this happens at a molecular level that has eluded resolution for half a century of intensive research. We studied the bearded dragon, a lizard that has sex chromosomes (ZZ male and ZW female), but in which that temperature can override ZZ sex chromosomes to cause male to female sex reversal. This provides an unparalleled opportunity to disentangle, in the same species, the biochemical pathways required to make a female by these two different routes. We sequenced the transcriptomes of gonads from developing ZZ reversed and normal ZW dragon embryos and discovered that different sets of genes are active in ovary development driven by genotype or temperature. Females whose sex was initiated by temperature showed a transcriptional profile consistent with the recently-proposed Calcium-Redox hypotheses of cellular temperature sensing. These findings are an important for understanding how the environment influences the development of sex, and more generally how the environment can epigenetically modify the action of genes.

## Introduction

Sex determination in vertebrates may be genetic or environmental. In genetic sex determination (GSD), offspring sex is determined by sex chromosomes inherited from each parent, which bear either a dominant gene on the heteromorphic sex chromosome (as with *SRY* in humans) (1,2), or a dosage sensitive gene on the homomorphic sex chromosome (as with *DMRT1* in birds) (3). However, some fish and many reptile species exhibit environmental sex determination (ESD), whereby a variety of external stimuli can determine sex, most commonly involving temperature (temperature dependent sex determination, TSD) (4,5). While GSD and ESD are commonly viewed as a dichotomy, the reality is far more complex. Sex determination in vertebrates exists as a continuum of genetic and environmental influences (6) whereby genes and environment can interact to determine sex (7–9).

The genetic mechanisms that act in highly conserved pathways that ultimately yield testes or ovaries are quite well characterised (5,10,11). Yet, despite decades of research on ESD systems, and TSD in particular, the upstream mechanisms by which an external signal is transduced to determine sex remains unknown (12). Recent research led to the hypothesis that the cellular sensor initiating ESD is controlled by the balance of redox regulation and calcium (Ca^2+^) signalling (CaRe) (13). The CaRe hypothesis proposes a link between CaRe sensitive cellular signalling and the highly conserved epigenetic processes that have been implicated in thermolabile sex (TSD and temperature sex reversal) (12,14–17). The CaRe hypothesis posits that in ESD systems a change in intracellular Ca^2+^ (probably mediated by thermosensitive transient receptor potential TRP channels) and increased reactive oxygen species (ROS) levels caused by high temperatures, alter the CaRe status of the cell, triggering cellular signalling cascades that drive differential sex-specific expression of genes to determine sex. The CaRe hypothesis makes several testable predictions for how an environmental signal is captured and transduced by the gonadal cells to deliver a male or a female phenotype.

Species in which genes and environment both influence sex determination provide unique opportunities to directly compare the regulatory and developmental processes involved in sex determination. By early gonad differentiation directed by genotype and temperature, it is possible to assess predictions of the CaRe hypothesis. In our model species, the central bearded dragon (*Pogona vitticeps*), we can compare female development via thermal and genetic cues because extreme temperatures (>32°C) override the male sex-determining signal from the ZZ sex micro-chromosomes to feminise embryos (8,18). This makes it possible to distinguish between the previously confounded effects of thermal stress and phenotypic sex by comparing gene expression throughout embryonic development in sex reversed ZZf females with genetic ZWf females.

We can explore the predictions of the CaRe model, namely that under sex-reversing conditions, we will see differential regulation of: 1) genes involved in responding to Ca^2+^ influx and signalling; 2) genes involved in antioxidant and/or oxidative stress responses; 3) genes with known thermosensitivity, such as heat shock proteins; 4) candidate TSD genes, such as *CIRBP* and Jumonji family genes; 5) signal transduction pathways such as the JAK-STAT and NF-ĸB pathways.

We compared gene expression profiles in *P. vitticeps* embryonic gonads at three developmental stages (6, 12 and 15; 19,20) for ZWf and ZZf eggs incubated at 28°C and 36°C respectively (Fig. 1). This allowed us to compare drivers of sex determination and differentiation under genetic or thermal influence. We found that very different regulatory processes are involved in temperature-driven regulation compared to gene-driven regulation, although both lead to a conserved outcome (ovaries, Fig. 2). We discovered dramatic changes in cellular calcium homeostasis in the gonads of ZZf individuals incubated at high sex reversing temperatures, which fulfill predictions of the CaRe hypothesis that this is the key driver of temperature induced feminization. We argue that differential expression of calcium channels, and subsequent alterations of the intracellular environment combined with increased ROS production encode, then transduce, the thermal signal into altered gene expression, ultimately triggering male to female sex reversal in *P. vitticeps*.

**Fig 1:**
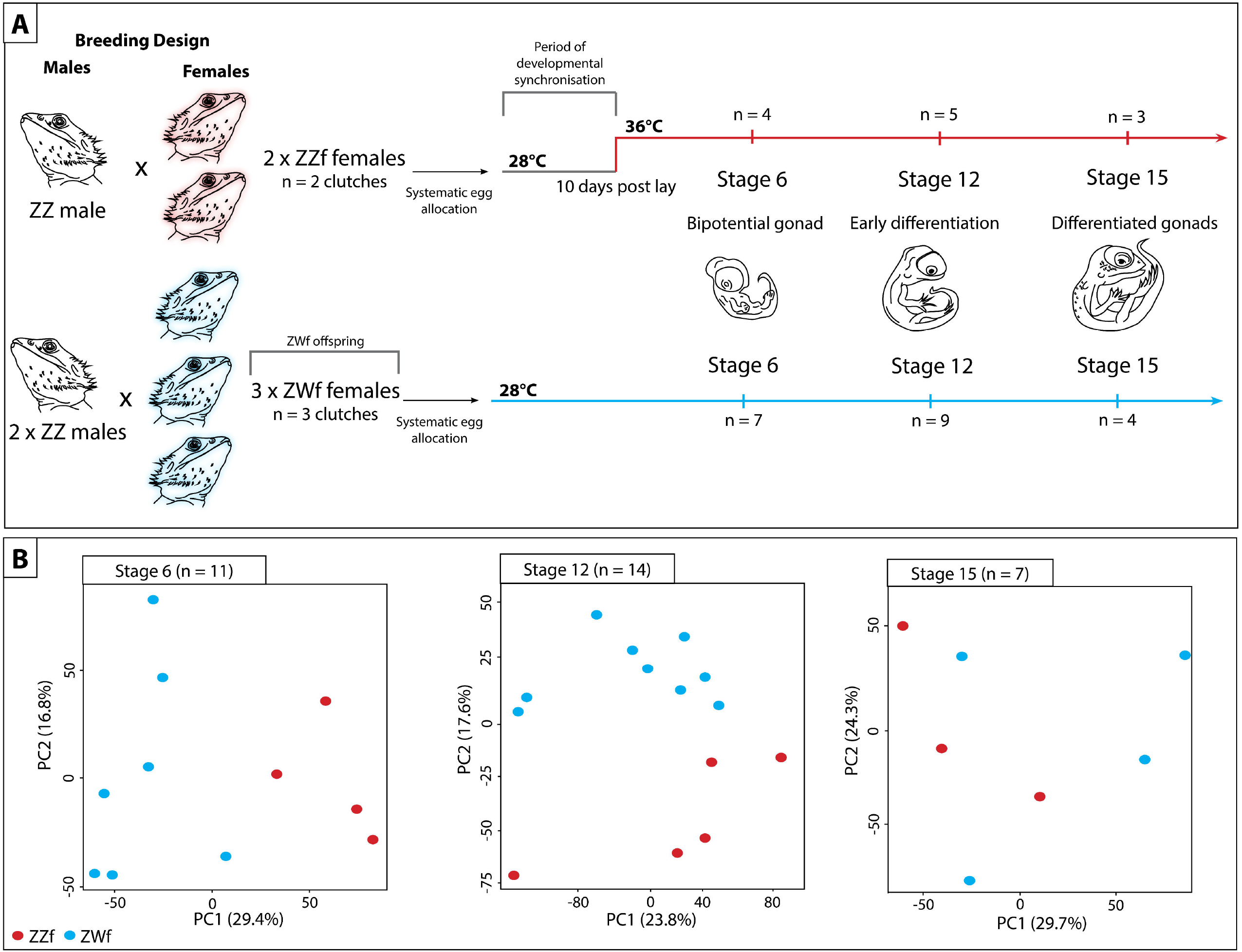
Schematic representation of experimental design used in this study to compare the differences between genetic sex determination and temperature dependent sex determination. (**A**) Summary of experiment showing how the parental crosses were designed, and how eggs were allocated and incubated. Eggs from sex reversed females (ZZf) were initially incubated at 28°C for 10 days, then were switched to 36°C. Eggs were sampled at the same three developmental stages (6, 12, and 15) based on (19,20). At stage 6 the gonad is bipotential, at stage 12 the gonad is in the early stages of differentiation, and it completely differentiated by stage 15. Eggs from concordant females (ZWf) were incubated at 28°C and sampled at the same three developmental stages as the ZZf eggs. (**B**) PCA plots showing the first and second principal components of read count per gene between ZZf (red) and ZWf (blue) at each stage of development.

**Fig 2:**
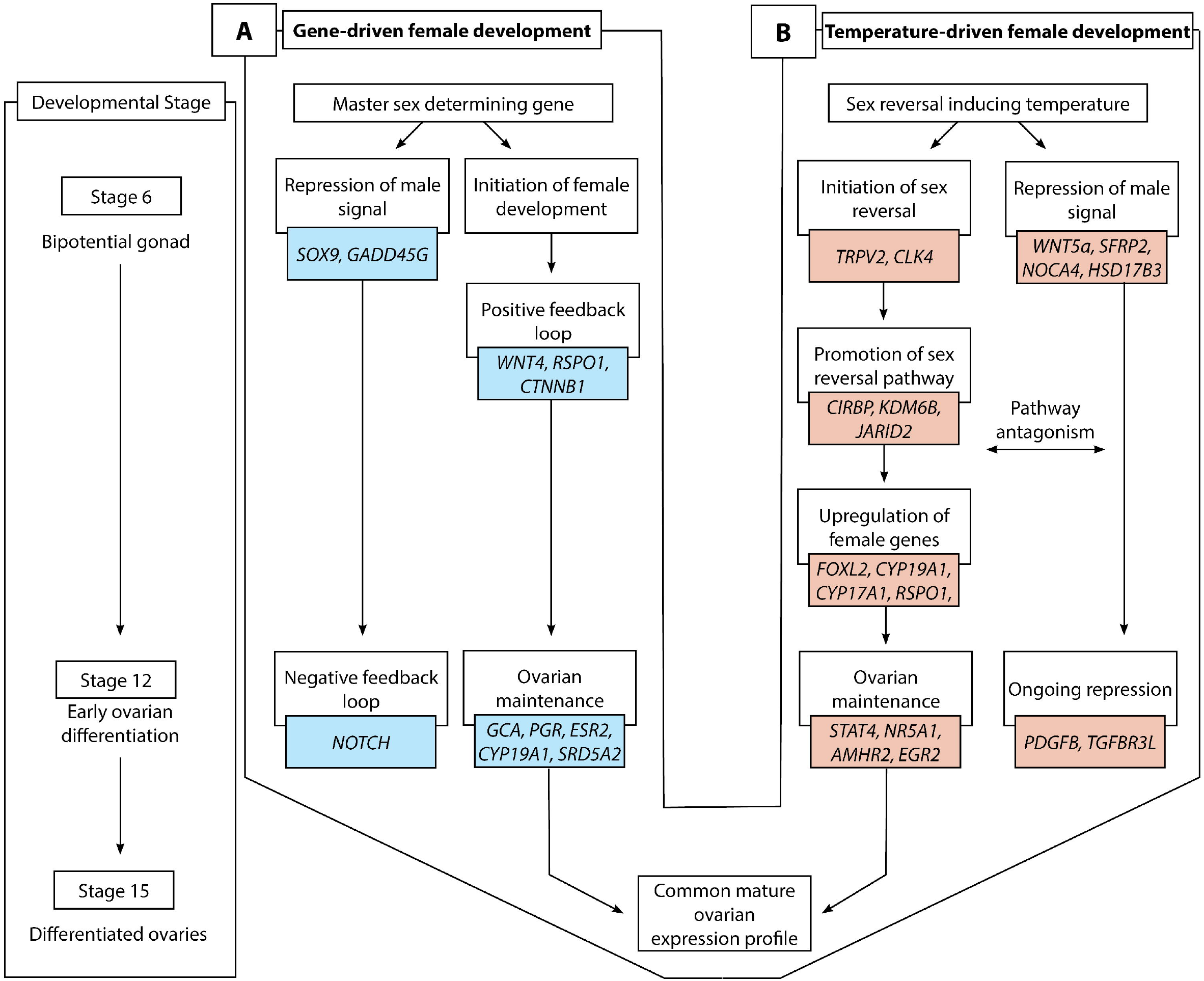
Schematic overview of gene-driven (blue) and temperature-driven (red) female developmental pathways in *Pogona vitticeps*. The pathways are initially different (from stages 6 to 12), but they ultimately converge on highly similar expression profiles when ovarian differentiation has occurred by stage 15. Both pathways are characterised by repression of a male signal, however this signal is stronger in temperature-driven females and appears to require ongoing repression when compared with the gene-driven females.

## Results

### Gene-driven female determination in ZWf embryos

Comparisons between stages in ZWf embryos (Fig. 3B, Fig. S1, Additional file S1) showed that many genes were differentially expressed between stages 6 and 12 (210 genes downregulated and 627 genes upregulated at stage 12), but few genes were differentially expressed between stages 12 and 15 (2 genes upregulated at stage 15).

**Fig 3:**
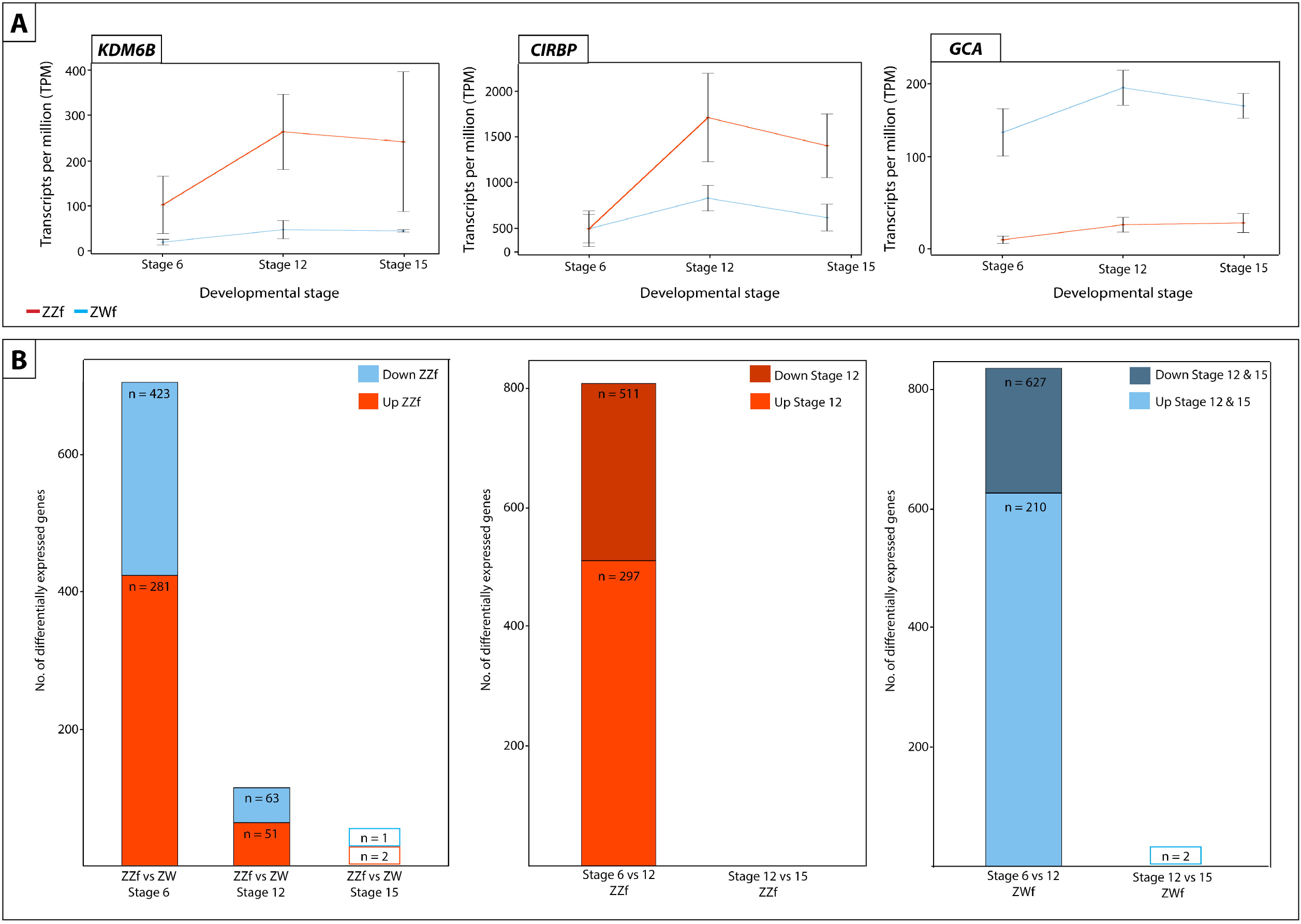
(**A**) Expression (transcripts per million, TPM) ± SE of three genes differentially expressed at all three developmental stages between ZZf and ZWf, with *KDM6B* and *CIRPB* (outlined in red) having consistently higher expression in ZZf embryos, and GCA having higher expression in ZWf. (**B**) Bar graphs representing the number of differentially expressed genes in all comparisons between ZZf and ZWf, and between developmental stages. MA plots of this data are available in Fig. S1. Differentially expressed genes were determined as having *P* values ≤ 0.01 and log_2_-fold changes of 1, −1.

*SOX9* and *GADD45G*, genes strongly associated with male development in mammals, were downregulated from stage 6 to stage 12, whereas various female related genes were upregulated, such as *PGR, ESR2, CYP19A1*, and *CYP17A1. BMP7*, a regulator of germ cell proliferation was upregulated at stage 12 (21), as were components of the NOTCH signalling pathway (*JAG2, DLL3, DLL4*), which are required for the suppression of Leydig cell differentiation (22,23). *SRD5A2*, whose product catalyses the 5-α reduction of steroid hormones such as testosterone and progesterone, was also upregulated (24,25).

Notably, there was little differential expression between stages 12 and 15, suggesting that genetically driven ovarian development is complete by stage 12 (Additional file S1).

### Temperature-driven female determination in ZZf embryos

Differential expression analysis of temperature-driven female development in ZZf embryos revealed many genes are differentially expressed between stages 6 and 12 (297 downregulated and 511 upregulated at stage 12) and no genes are differentially expressed between stage 12 and 15 (Fig. 3, Fig. S1, Additional file S1), suggesting completion of the ovarian development by stage 12 also in ZZf females.

Upregulation of *FZD1*, a receptor for *Wnt* family proteins required for female development, suggests the activity of female pathways in ZZf embryos (26). As was seen for ZWf females, canonical NOTCH ligands *DLL3* and *DLL4* were upregulated from stage 6 to stage 12 in ZZf females. However, this did not coincide with upregulation of JAG ligands or NOTCH genes, and the GO term “negative regulation of NOTCH signalling” was enriched within the group of genes upregulated from stage 6 to 12 in ZZf females (Additional file S2). Further, *PDGFB*, which is required for Leydig cell differentiation, was upregulated (27).

Together, this suggests that the NOTCH signalling pathway may not be activated, and Leydig cell recruitment is not strongly repressed at stage 12 in ZZf. Alternatively, the absence of NOTCH signalling may indicate an important transition from progenitor cells to differentiated gonadal cell types in the early stages of the developing ovary (28). These apparent differences in NOTCH signalling between ZZf and ZWf embryos suggests that ovarian development has progressed further in ZWf females.

Interestingly, genes typically associated with male development show diverse regulation in ZZf embryos, with some being downregulated and some being upregulated from stage 6 to 12. These included *WNT5a* and *SFRP2*, which are both involved in testicular development in mice (29,30), *NCOA4*, which enhances activity of various hormone receptors, and exhibits high expression in testes in mice during development (31), and *HSD17B3*, which catalyses androstenedione to testosterone (32). Unlike what was observed in ZWf embryos, *SOX9* and *GADD45G* were not differentially expressed between stages 6 and 12 in ZZf embryos. *TGFBR3L*, which is required for Leydig cell function in mouse testis (33), and *NR5A1, SOX4*, and *AMHR2* (34–37) were also differentially expressed between stages 6 and 12 (Additional file S1).

A suite of genes typically associated with female development were upregulated from stage 6 to 12 (38), for example, *FOXL2, CYP17A1, RSPO1*, and *ESRRG*. As was also observed in stage 12 ZWfs, *ESR2, BMP7, CYP19A1*, and *PGR*, were more highly expressed at stage 12 in ZZfs. Notably, *CYP19A1* was much more strongly upregulated at stage 12 in ZZfs compared with stage 12 in ZWfs (Additional file S1). The increase in sex specific genes was also reflected in enriched GO terms at stage 12, which included “hormone binding”, “steroid hormone receptor activity”, and “female sex determination” (Additional file S2).

### Ovarian maintenance in sex reversed ZZf females

The maintenance of female gene expression and ovarian development at stage 12 in ZZf females may be centrally mediated by *STAT4* (Fig. 4). As a member of the *JAK-STAT* pathway, *STAT4* is transduced by various signals, including reactive oxygen species, to undergo phosphorylation and translocate from the cytoplasm to nucleus (39–41). At stage 12, *STAT4* is upregulated, alongside *PDGFB* compared to stage 6 in ZZf females. *PDGFB* is known to activate *STAT4* (40). Various *STAT4* target genes, notably *AMHR2, NR5A1, EGR1*, and *KDM6B* (40) are also upregulated at stage 12 (Additional file S1). Consistent with this link is the observation that a member of the same gene family, *STAT3*, is implicated in TSD in *Trachemys scripta* (17).

**Fig 4:**
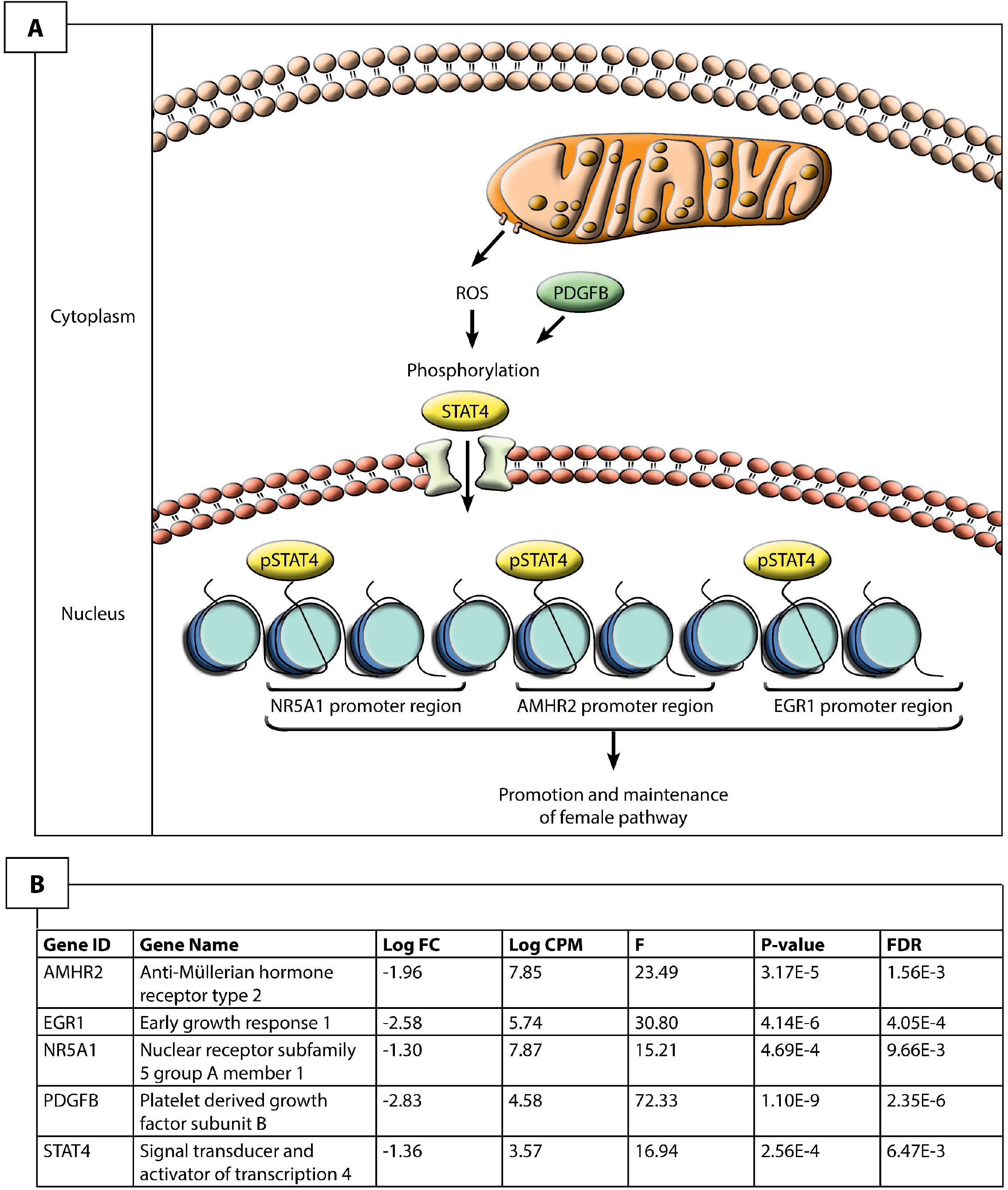
Hypothesized pathway for the maintenance of the ovarian phenotype in stage 12 sex reversed ZZf *Pogona vitticeps*. Given the upregulation of these genes, it is likely that reactive oxygen species induce the phosphorylation, and subsequent activation and nuclear translocation of STAT4, likely mediated by PDGFB. Once in the nucleus, STAT4 is able to bind to promoter regions of known target genes, *NR5A1, AMHR2* and *EGR2* to regulate their expression and promote ovarian development.

Several targets of *STAT4* are upregulated at stage 12, including *AMHR2* and *NR5A1*. Though typically associated with male development, *AMHR2* and *NR5A1* may also have roles in ovarian development. Although it is the primary receptor for *AMH, AMHR2* exhibits considerable evolutionary flexibility is sometimes associated with ovarian development (reviewed by (36)). *NR5A1* is also often associated with male development, as it positively regulates expression of *AMH* and *SOX9* in mammals (42). However, *NR5A1* can also interact with *FOXL2* and bind to *CYP19A1* promoter to promote female development (43,44). The upregulation of *FOXL2* and *CYP19A1*, but not *AMH* or *SOX9*, suggests that *NR5A1* is involved in the establishment of the ovarian pathway in ZZf females.

*EGR1* positively regulates *DMRT1* expression through promoter binding in Sertoli cells, but knock-out of this gene can also cause female infertility in mice (45–47). EGR1 is also associated with female development in birds, likely controlling the production of steroid hormones (34). As was observed for *NR5A1, DMRT1* was also lowly expressed, suggesting it is not activated by *EGR1* in ZZf females.

One explanation for these expression trends is that male-associated genes are not strongly repressed at this stage during the sex reversal process, and that more prolonged exposure to the sex reversing temperature is required to firmly establish the female phenotype. However, we argue that the results more strongly suggest ROS-induced activation of *STAT4*, and subsequent phosphorylation and translocation, probably mediated by *PDGFB*, allows for the transcriptional activation of *NR5A1, AMHR2*, and *EGR1*, which in the temperature driven process of sex reversal in the ZZf embryos serve to maintain the ovarian phenotype.

### Differential regulation of female developmental pathways

To better understand differences in ovarian developmental pathways, we compared gene expression of ZZf with ZWf embryos at each developmental stage. There are large gene expression differences between normal ZWf females and ZZf sex reversed females early in development, before the bipotential gonad differentiates into an ovary. These differences are most pronounced early in development and diminish as development progresses. Stage 6 had the largest number of differentially expressed genes (DEGs) (281 genes higher expressed in ZWf embryos, 423 genes higher expressed in ZZf), with fewer DEGs at stage 12 (51 genes upregulated in ZWf, 63 genes upregulated in ZZf), and fewest at stage 15 (1 gene upregulated in ZWf, 2 genes upregulated in ZZf) (Fig. 3B, Additional file S3, Fig. S1). This suggests that the sex reversed embryos start out on a male developmental trajectory, which they pursue beyond the thermal cue (3 days when the eggs were switched to high incubation temperatures, Fig. 1), but by stage 12 development has been taken over by female genes.

Gene ontology (GO) enrichment analysis showed important differences between ZZf and ZWf at stage 6, and provides independent support for the role of calcium and redox regulation in ZZf females as proposed by the CaRe model (Fig. 5 A-D, Additional file S4). GO processes enriched in the gene set higher expressed in ZZf at stage 6 included “oxidation-reduction processes”, “cytosolic calcium ion transport”, and “cellular homeostasis” (Fig. 5A, Additional file S4). GO function enrichment also included several terms related to oxidoreductase activities, as well as “active transmembrane transporter activity” (Fig. 5C, Additional file S4). No such GO terms were enriched in the gene set higher expressed in ZWf. Instead, enriched GO terms included “anatomical structure development”, and “positive regulation of developmental growth” (Fig. 5B, D, Additional file S4).

**Fig 5:**
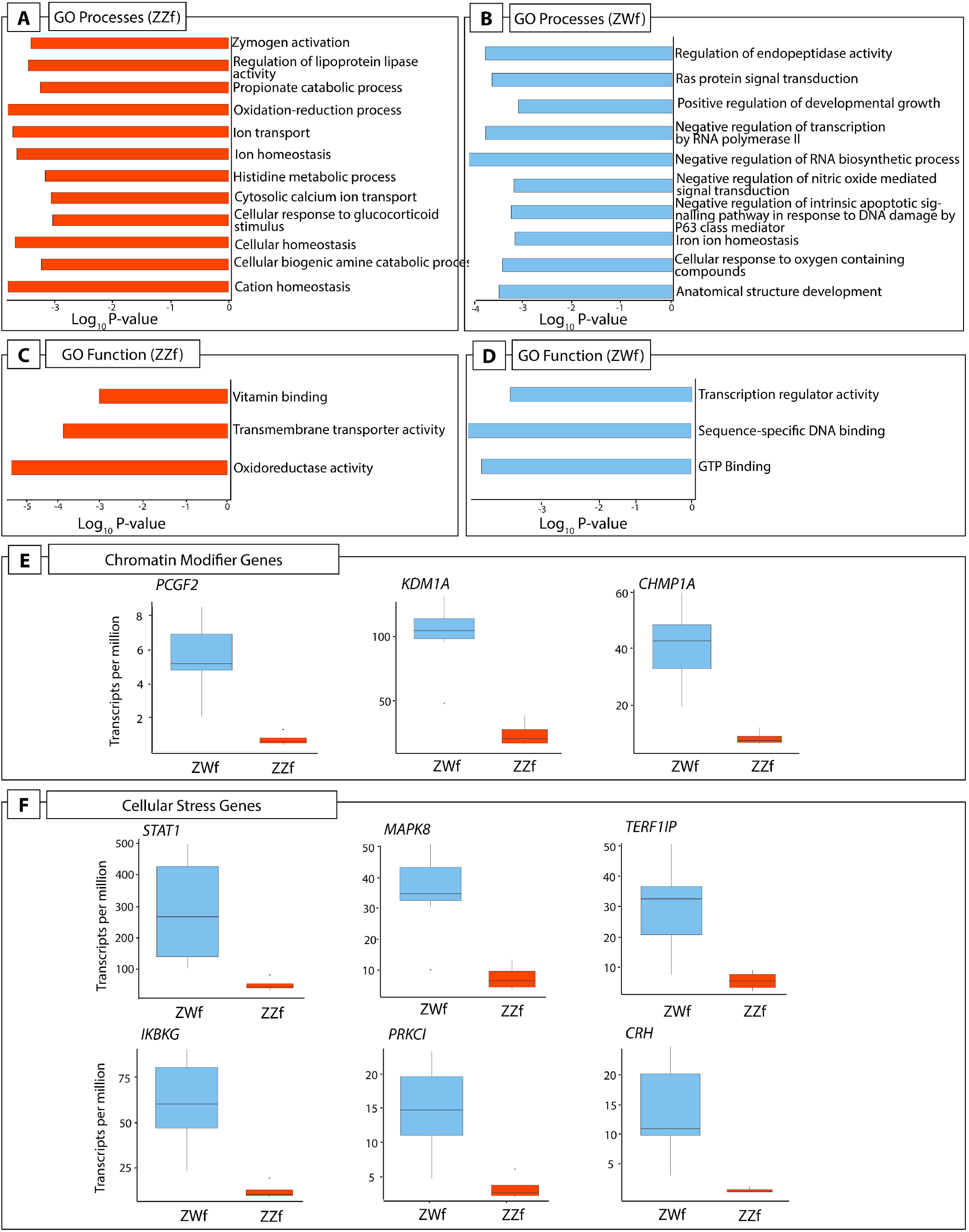
(**A**) A subset of GO processes and (**C**) GO functions enriched in stage 6 ZZf embryos compared with ZWf. (**B**) A subset of GO processes and (**D**) GO functions enriched in stage 6 ZWf embryos compared with ZZf. Complete results of GO analysis for all developmental stages in ZZf and ZWf for enriched GO processes and functions is provided in Additional file S2. Differentially expressed chromatin modifier (**E**) and cellular stress (**F**) genes in *Pogona vitticeps* at stage 6 comparing ZWf and ZZf females.

Genes involved in female sex differentiation were higher expressed at stage 6 in normal ZWf embryos compared to sex reversed ZZf embryos (Additional file S3). These included *FOXL2, ESR2, PGR*, and *GATA6* (48,49). Higher expression of *LHX9*, a gene with a role in bipotential gonad formation in mammals and birds, was more highly expressed in ZWf embryos (42,50–52). Two genes with well described roles in male development, *SOX4* and *ALDH1A2* (53–55), were also higher expressed in ZWf embryos, suggesting they have an as yet unknown function in the early establishment of the ovarian trajectory in *P. vitticeps*.

Taken together, these results further suggest that ZWf females are committed to the female pathway earlier than ZZf females. This is not surprising, since ZWf females possess sex chromosomes from fertilisation, whereas ZZf individuals have had only 3 days of exposure to a sex reversal inducing incubation temperature (Fig. 1A). This data is the first to demonstrate a difference in timing of genetic signals between gene and temperature driven development in the same species.

Three genes were constantly differentially expressed between ZWf and ZZf embryos at all three developmental stages (Fig. 3A). *GCA* (grancalcin) was upregulated in ZWf embryos, and *KDM6B* and *CIRBP* were upregulated in ZZf embryos at all developmental stages. GCA is a calcium binding protein commonly found in neutrophils and is associated with the Nf-κB pathway (56,57). It has no known roles in sex determination, but its consistent upregulation in ZWf embryos compared to ZZf embryos suggests GCA is associated with gene driven ovarian development, at least in *P. vitticeps*.

Further analysis of gene expression trends using K-means clustering analysis (58) was used to investigate genes associated with female development, and to determine to what extent these genes are shared between ZZf and ZWf embryos (Fig. 6, Additional file S5). Clusters with upward trends reflect genes likely to be associated with female development, so clusters 1 and 4 in ZWf (ZWC1 and ZWC4), and clusters 1 and 2 in ZZf (ZZC1 and ZZC2), were explored in greater detail (Fig. 6, Additional file S5).

**Fig 6:**
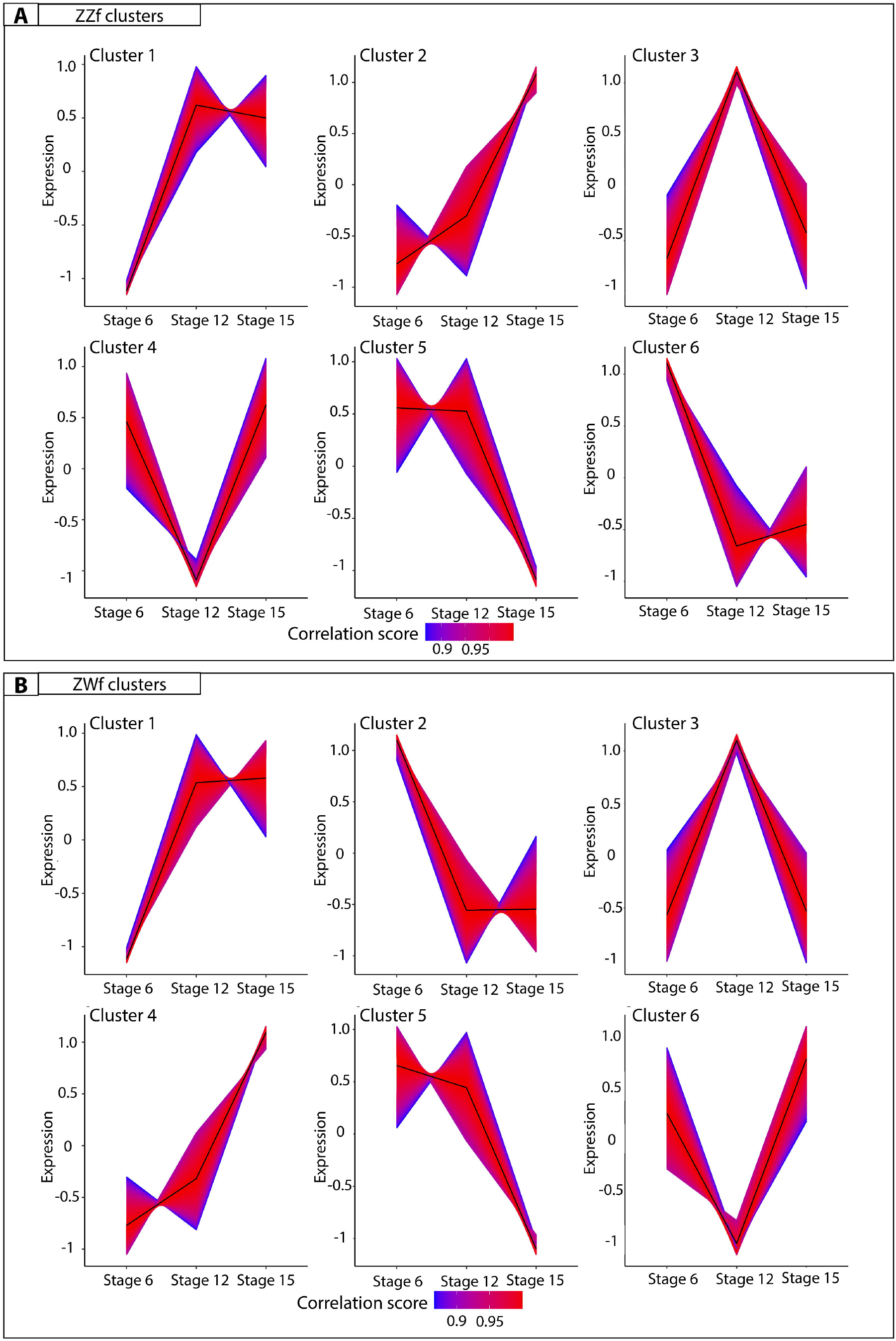
K-means clustering analysis on normalized counts per million for ZZf (**A**) and ZWf (**B**) across all developmental stages. The colour depicts the correlation score of each gene in the cluster, where numbers approaching one (red) have the strongest correlation. All gene lists produced for each cluster are provided in Additional file S5.

ZWC4 and ZZC2 shared 374 genes. Enriched GO terms included “germ cell development” and “reproductive processes” (Additional file S6), consistent with a link with female development. Genes identified included *FIGLA*, a gene known to regulate oocyte-specific genes in the female mammalian sex determination pathway (59), and *STRA8* which controls entry of oocytes into meiosis. Intriguingly, the GO term “spermatid development” was also enriched, encompassing many genes with known roles in testes function, including *ADAD1* and *UBE2J1* (60,61). This suggests that genes involved in male sex determination in mammals may have been co-opted for use in the ovarian pathway in reptiles, so their roles require further investigation in other vertebrate groups, particularly given the complex nature of gene expression in sperm cell types.

ZWC1 and ZZC1 shared 998 genes. ZZC1 has about 700 unique genes and ZWC1 about 500. GO analysis on shared genes between these clusters (n = 998) revealed enrichment terms such as “kinase binding” and “intracellular signal transduction” (Additional file S5). Genes unique to ZZC1 included members of heat shock protein families (*HSPB11, HSPA4, HSP90AB1, HSPH1, HSPB1, HSPD1*), heterogenous ribonucleoprotein particles (*HNRNPUL1*), mitogen activated proteins (including *MAPK1, MAPK9, MAP3K8*), and chromatin remodelling genes (*KDM2B, KDM1A, KDM5B, KDM3B*). GO enrichment for genes unique to ZZC1 included “mitochondrion organisation”, “cellular localisation”, and “ion binding”, while GO enrichment for genes unique to ZWC1 included “regulation of hormone levels” and numerous signalling related functions (Additional file S6).

Taken together, our results show that although the same ovarian phenotype is produced in genetic and temperature induced females, this end is achieved via different gene expression networks. This is most pronounced at stage 6, after which the extent of the differences decreases through development. This reflects canalisation of the gonadal fate to a shared outcome (ovaries, Fig. 2).

### Signature of hormonal and cellular stress in ZWf females

Previous work on *P. vitticeps* has shown a more than 50-fold upregulation of a hormonal stress response gene, *POMC*, in sex reversed adult females, leading to the suggestion that induction of sex reversal is in response to temperature stress, or that it is an inherently stressful event, the effects of which persist into adult life (14). We therefore investigated the expression of stress related genes in ZZf and ZWf embryos.

We found considerable evidence that ZZf embryos experience oxidative stress, likely resulting from increased ROS production (discussed in detail below). However, contrary to our expectations, we found that ZWf embryos showed higher expression than ZZf of hormonal stress genes and pathways that have been hypothesized to be involved in sex reversal (Fig. 5E-F, Additional file S3). Genes upregulated in ZWf embryos compared to ZZf embryos included *STAT1*, a component of the JAK-STAT pathway, with several roles in stress responses (62), and *MAP3K1* and *MAPK8*, which are typically involved in mediating stress-related signal transduction cascades (63–65). *TERF2IP* is also upregulated; this gene is involved in telomere length maintenance and transcription regulation (66). When cytoplasmic, *TERF2IP* associates with the l-kappa-B-kinase (IKK) complex and promotes IKK-mediated phosphorylation of RELA/p65, activating the NF-κB pathway and increasing expression of its target genes (67). Notably two members of the IKK complex, *IKBKG* (also known as *NEMO*) and *PRKCI*, which are involved in NF-κB induction, were also upregulated in ZWf embryos compared to ZZf embryos (Fig. 4b), implying activation of the NF-κB pathway (68). This pathway is typically associated with transducing external environmental signals to a cellular response (69,70), but also has diverse roles in sex determination in mammals, fish, and invertebrate models (reviewed by Castelli *et al*., 2019).

*CRH*, another gene upregulated at stage 6 in ZWf females compared with ZZf females (Fig. 5b, supplementary file S3), is best known for its role as a neuropeptide synthesised in the brain in response to stresses that trigger the hypothalamic-pituitary-adrenal (HPA) axis (71,72). The role of *CRH* production in the gonads, particularly in ovaries is currently poorly understood (73–76). High *CRH* expression in ZWf gonads is the first observation of this in reptiles. The role of the hormonal stress response during embryonic development, and its apparent discordance with results observed in adults in *P. vitticeps* requires further investigation (14).

### Cellular signalling cascades driving sex reversal

Results of this study provide considerable corroborative support for the CaRe model, which proposes a central role for calcium and redox in sensing and transducing environmental signals to determine sex. Many of the genes and pathways predicted by the CaRe model to be involved in sex reversal were shown to be upregulated in ZZf embryos at stage 6 compared to ZWf embryos. We use the CaRe model as a framework to understand the roles of each signalling participant in their cellular context during the initiation of sex reversal (Fig. 7). This interpretation is also independently supported by GO analysis, showing enrichment of expected terms, such as “cytosolic calcium ion transport” (Additional file S4), as well as k-means clustering analysis (Additional files S4, S5).

**Fig 7:**
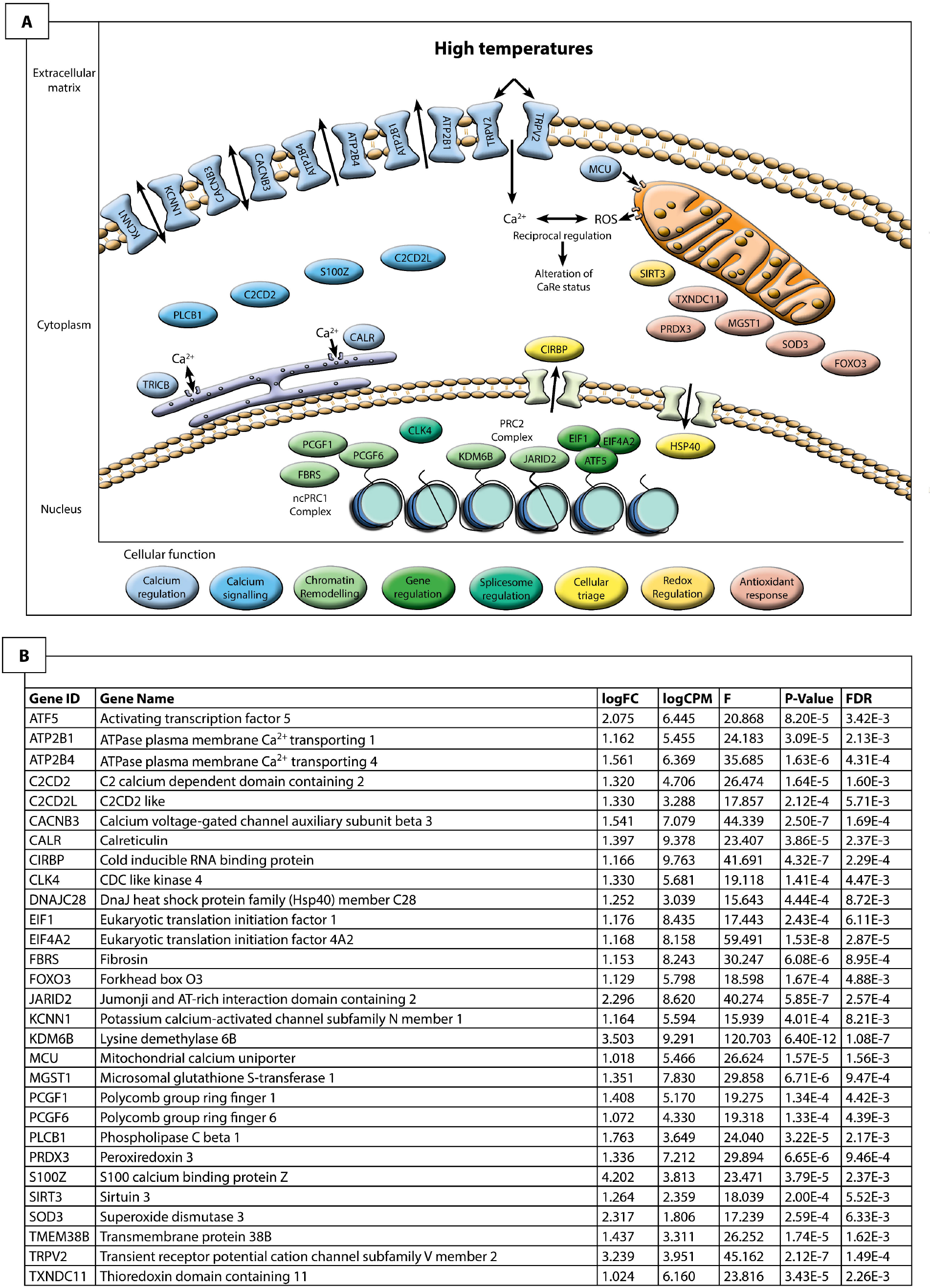
Hypothesised cellular environment (*A*) of a ZZf gonad at stage 6 in *Pogona vitticeps* based on differential expression analysis (*B*) using the CaRe model as a framework (13). We used this approach to understand the cellular context responsible for driving sex reversal in ZZf samples. This reveals that calcium signalling likely plays a very important role in mediating the temperature signal to determine sex. Influx of intracellular calcium is likely mediated primarily by TRPV2, and may also be influenced by KCNN1 and CACNB3. This influx appears to trigger significant changes in the cell to maintain calcium homeostasis. MCU, ATP2B1, CALR and TRICB all play a role in this process by sequestering calcium and pumping it back out of the cell, in which KCNN1 and CACNB3 may have a role. Calcium signalling molecules C2CD2, C2CDL2, and S100Z are likely responsible for encoding and translating the calcium signal leading to changes in gene transcription. Changes in gene expression are likely mediated primarily by the two major Polycomb Repressive Complexes, PRC1 and PRC2. Members of these two complexes (PCGF1, PCGF6, KDM6B, and JARID2) transcriptionally regulate genes by controlling methylation dynamics of their targets, the latter two of which have been previously implicated in sex reversal (14,15). ATF5 may also play a role in gene regulation, and alternative splicing, which has been implicated in sex reversal (14) may be mediated by CLK4. High temperatures necessarily increase cellular metabolism, which in turn increases the amount of reactive oxygen species (ROS) produced by the mitochondria. ROS can cause cellular damage at high levels, so trigger an antioxidant response, which is observed here in the upregulation of MGST1, PRDX3, TXNDC11 and FOXO3. Also of note is the upregulation of CIRBP, which has numerous functions in response to diverse cellular stresses, and has been implicated in TSD.

Cluster 6 in ZZf (Fig. 6A) shows genes whose expression decreases after stage 6, so is likely to include genes responsible for the initial response to temperature and initiation of sex reversal, whose continuing action is not required once the ovarian trajectory has been established. Consistent with this assumption, as well as with predictions from the CaRe model, the 4050 genes in this cluster were enriched for GO terms that included “oxidation-reduction process” and various oxidoreductase activities (Additional files S5, S6).

### Calcium transport, signalling, and homeostasis

Our data suggest that exposure to high temperatures may cause a rapid increase in cytosolic Ca^2+^ concentrations, as calcium influx is probably mediated by the thermosensitive calcium channel, *TRPV2* (77,78). *TRPV2* was upregulated in stage 6 ZZf embryos compared to ZWf embryos (Fig. 6). Transient receptor potential (TRP) ion channels, including *TRPV2*, have previously been implicated in TSD in *Alligator* sinensis and *A. mississippiensis*, as well as the turtle *Trachemys scripta* (79–82).

*TRPV2* mediated Ca^2+^ influx may trigger a cascade of changes within the gonadal cells of ZZf females, which restore calcium homeostasis, critical to avoid apoptosis (83,84). We observed evidence of such a homeostatic response, with the upregulation of seven genes involved in Ca^2+^ transport and sequestration in ZZf females compared to ZWf females at stage 6 (Fig. 6). Specifically, *MCU, ATP2B1, ATP2B4*, together regulate calcium homeostasis through active transport of calcium into the mitochondria and into the extracellular space (85–87). *KCNN1* and *CACNB3* encode proteins required for the formation of plasma membrane channels controlling the passage of Ca^2+^ (88–90). *CACNB3* and *KCNN1* have well characterised roles in the nervous system, and excitable cell types in muscle, but their association with TSD in embryonic gonads is novel (88,89). Evidence is also building for a broader role for voltage-gated calcium channels, including *CACNB3*, in orchestrating Ca^2+^ signalling and gene regulation (91). We suggest that *KCNN1* and *CACNB3* in gonads of TSD species play roles in mediating the homeostatic response to elevated cytosolic Ca^2+^ concentrations, and are involved in the subsequent modulation of Ca^2+^ signalling pathways.

*TRPC4*, another TRP family gene was, upregulated in stage 12 compared to stage 6 in both ZZf and ZWf embryos. *TRPC4* is expressed in mouse sperm and inhibited by progesterone (92,93) but has no known association with sex determination. *TRPC4* belongs to the TRPC superfamily, which all conduct calcium ions into the cell, typically through phospholipase C and calmodulin signalling pathways, G-protein-coupled receptors, and receptor tyrosine kinases (94,95). Notably, *PLCL2* a phospholipase gene, together with calmodulin genes *CALM1* and *CAMKK1*, were upregulated alongside *TRPC4* from stage 6 to stage 12 in ZZf embryos but not in ZWf embryos (Additional file S3). Given *TRPC4* is upregulated from stage 6 to 12 in both ZZf and ZWf females, it may play a more conserved role in ovarian development in *P. vitticeps*.

Several genes with functions in calcium metabolism were upregulated in stage 6 ZZf embryos compared to stage 6 ZWf embryos. *CALR* encodes a multifunctional protein that acts as a calcium binding storage protein in the lumen of the endoplasmic reticulum, so is also important for regulating Ca^2+^ homeostasis (84,96,97). *CALR* is also present in the nucleus, where it may play a role in regulation of transcription factors, notably by interacting with DNA-binding domains of glucocorticoid and hormone receptors, inhibiting the action of androgens and retinoic acid (97–101). *TMEM38B* (commonly known as *TRICB*) is also found on the endoplasmic and sarcoplasmic reticula, where it is responsible for regulating the release of Ca^2+^ stores in response to changes in intracellular conditions (102).

*MCU, ATP2B1, ATP2B4, KCNN1, CACNB3, CALR*, and *TMEM38B* have no known roles in vertebrate sex determination, so their association with sex reversal in *P. vitticeps* is new. This upregulation during the early stage of sex reversal suggests that they are upstream modulators involved in the transduction of environmental cues that trigger sex determination cascades, which is consistent with predictions made by the CaRe hypothesis.

We hypothesize that intracellular Ca^2+^ increases in stage 6 ZZf gonads, and further observe that Ca^2+^ signalling related genes are also upregulated in ZZf females compared to stage 6 ZWf females (Fig. 7). *C2CD2* and *C2CD2L* are both thought to be involved in Ca^2+^ signalling, although there is no functional information about these genes. Of note is the significant upregulation of *S100Z*, which is a member of a large group of EF-hand Ca^2+^ binding proteins that play a role in mediating Ca^2+^ signalling (103). The EF-hand domain is responsible for binding Ca^2+^, allowing proteins like that encoded by *S100Z* to ‘decode’ the Ca^2+^ biochemical signal and translate this to various targets involved in many cellular functions including Ca^2+^ buffering, transport, and enzyme activation (104,105). *PLCB1* also contains an EF-hand binding domain and behaves similarly, being activated by many extracellular stimuli and effecting numerous signalling cascades. It can translocate to the plasma membrane and nucleus, and release Ca^2+^ from intracellular stores (106). Some Ca^2+^ related genes (*GCA* and *CALM1*) are also upregulated in ZWf embryos, but make only a small proportion of the overall response in differential gene expression (Additional file S3).

### Oxidative stress in response to high temperatures

The upregulation of antioxidant genes in ZZf compared to ZWf embryos suggests that the gonadal cells in the ZZf embryos are in a state of oxidative stress (Fig. 7). As was proposed by the CaRe model, we see results consistent with the prediction that high incubation temperatures increase metabolism, which increases the production of reactive oxygen species (ROS) by the mitochondria, resulting in oxidative stress (13). ROS are required for proper cellular function, but above an optimal threshold, they can cause cellular damage (107,108). Crossing this threshold launches the antioxidant response, which causes the upregulation of antioxidant genes to produce protein products capable of neutralising ROS (109,110). We observed upregulation of redox related genes, specifically of *TXNDC11, PRDX3, MGST1* in ZZf embryos compared to ZWf embryos at stage 6. Also upregulated was *FOXO3*, which plays a role in oxidative stress responses, typically by mediating pro-apoptotic cascades (111,112). Importantly, antioxidants play other cellular roles besides neutralisation of ROS. One of these is the alteration of cysteine resides through a process known as S-glutathionylation (113).

Various redox related genes were downregulated from stage 6 to 12 in ZZf embryos but not in ZW embryos, including *GLRX* and *PRDX3* (114), as well as numerous genes involved in ROS induced DNA damage repair; *LIG4, ENDOD1*, and *HERC2* (115). This indicates a need for expression of these genes specifically in ZZf embryos in early stages that ceases in transition to stage 12. *STAT4*, a member of the ROS-induced JAK-STAT pathway (Simon et al. 1998), and *DDIT4*, which is involved stress responses to DNA damage (116), were both upregulated from stage 6 to stage 12 in ZZf embryos.

The vertebrate antioxidant response is typically initiated by *NRF2*, but we observed no differential expression of *NRF2*, only upregulation of some of its known targets in ZZf embryos (117). This may mean that the action of *NRF2* is depends more on its translocation from the cytoplasm to the nucleus to modulate transcription of target genes, a process that does not necessarily rely on increased expression of *NRF2* (117). Alternatively, NRF2 upregulation may have occurred prior to sampling.

Oxidative stress has previously been proposed to have a role in TSD, based on the upregulation of genes involved in oxidative stress response. One of these genes, *UCP2*, was upregulated at high male producing temperatures in *A. mississippiensis* (82). UCP2, and others genes involved in oxidative stress responses, were also implicated in UV induced masculinisation in larvae of a thermosensitive fish species (*Chirostoma estor*) (118). Notably, we found that *UCP2* was upregulated between stages 6 to 12 in ZZf *P. vitticeps* embryos, suggesting a sustained response to thermal stress in the mitochondria (Additional file S3).

### Temperature response and cellular triage

We also observed upregulation of genes involved in response to more generalised environmental stress in ZZf compared to ZWf embryos, as expected since the embryos exposed to high temperature were experiencing a state of thermal stress (Fig. 7). Notably, *CIRBP* a promising candidate for regulation of sex determination under thermal influence, is approximately 10-fold upregulated in ZZf compared to ZWf (Additional file S3). *CIRBP* has a highly conserved role in generalised stress responses (119). It has been suggested to be a putative sex determining gene in the TSD turtle *Chelydra serpentina* (120), and is differentially expressed at different incubation temperatures in *Alligator sinensis* (81). We also observed the upregulation of *CLK4* in ZZf compared to ZWf embryos, a gene that has been recently shown to be inherently thermosensitive, and to regulate splicing of temperature specific *CIRBP* isoforms (121).

We found that *ATF5* is upregulated in ZZf embryos compared to ZWf embryos (Fig. 7). *ATF5* has diverse roles in stimulating gene expression or repression through binding of DNA regulatory elements. It is broadly involved in cell specific regulation of proliferation and differentiation, and may also be critical for activating the mitochondrial unfolded protein response (122). This gene is induced in response to various external stressors, and is activated via phosphorylation by eukaryotic translation initiation factors, two of which (*EIF1* and *EIF4A2*; Zhou et al. 2008) are also upregulated in ZZf embryos compared to ZWf embryos.

Though not well studied in the context of sex determination, heat shock factors and proteins have been implicated in female sex determination in mammals and fish, and may also play a conserved role in the ovarian pathway in *P. vitticeps* (79,124–127). Surprisingly, only one gene associated with canonical heat shock response (*HSP40*, also known as *DNAJC28*) was differentially expressed following exposure to high temperature in stage 6 ZZf females compared to ZWf embryos (Additional file S3). This could mean either that a heat shock response occurs prior to sampling, or that *P. vitticeps* uses different mechanisms to cope with heat shock.

### Chromatin remodelling

We observed upregulation of several components of two major chromatin remodelling complexes, polycomb repressive complexes PRC1 and PRC2, in both the genotype-directed ZWf and the temperature-directed ZZf female pathways in *P. vitticeps* (Fig. 7). Chromatin modifier genes *KDM6B* and *JARID2* are involved in regulation of gene expression during embryonic development and epigenetic modifications in response to environmental stimulus (128,129). *JARID2* and *KDM6B* were both upregulated in ZZf embryos compared to ZWf embryos in stages 6 and 12, and *KDM6B* was also upregulated at stage 15. These genes have recently been implicated in two TSD species (*Alligator mississippiensis*, and *Trachemys scripta*) and temperature sex reversed adult *Pogona vitticeps* (14,15,79,82).

We also found that two other members of the PRC1 complex, *PCGF6* and *PCGF1*, were upregulated in ZZf embryos at stage 6 compared to ZWf embryos (Fig. 7). *PCGF6* is part of the non-canonical PRC1 complex (ncPRC1) that mediates histone H2A mono-ubiquitination at K119 (H2AK119ub) (130,131). *PCGF6* acts a master regulator for maintaining stem cell identity during embryonic development (132), and is known to bind to promoters of germ cell genes in developing mice (130). *PCFF1* exhibits similar functions by ensuring the proper differentiation of embryonic stem cells (133). The ncPRC1 complex also promotes downstream recruitment of PRC2 and H3K27me3, so that complex synergistic interactions between PRC1 (both canonical and non-canonical) and PRC2 can occur (134,135).

We found that other components of both PRC1 and PRC2 complexes were also upregulated in ZWf embryos compared to ZZf embryos (Fig. 5a, supplementary file S3). A member of the canonical PRC1 complex, *PCGF2* (also known as *MEL18*), was upregulated in ZWf embryos compared to ZZf embryos (134). This gene has previously been implicated in temperature induced male development in *Dicentrachus labrax* (136), and is required for coordinating the timing of sexual differential in female primordial germ cells in mammals (137). *KDM1A*, a histone demethylase that is required for balancing cell differentiation and self-renewal (138), was upregulated in ZWf embryos compared to ZZf embryos. *CHMP1A* was upregulated in ZWf, and is likely to be involved in chromosome condensation, as well as targeting PcG proteins to regions with condensed chromatin (139).

Thus, we conclude that the initiation of sex reversal in ZZf *P. vitticeps* involves a complex cascade of cellular changes initiated by temperature. Our data are consistent with the predictions of the CaRe hypothesis that high temperatures are sensed by the cell via TRP channels, which causes an increase in intracellular increase of Ca^2+^. Coincident with this is an increase of ROS production in the mitochondria that causes a state of oxidative stress. Together, Ca^2+^ and ROS alter the CaRe status of the cell, trigger a suite of alternations in gene expression including chromatin remodelling, which drives sex reversal (Fig. 7).

## Discussion

We used the unique sex characteristics of our model reptile species, *Pogona vitticeps*, which determines sex genetically but sex reverses at high temperature, to assess predictions of the CaRe hypothesis (13). By sequencing isolated embryonic gonads, we provide the first data to represent a suite of key developmental stages with comparable tissue types, and will be a valuable resource for this reptilian model system. There are few transcriptomes of GSD reptiles during embryonic development; the only dataset available prior to this study was a preliminary study of the spiny softshell turtle, *Apalone spinifera* (126), which was inadequate for the inter-stage comparisons required to explore genetic drivers of gonad differentiation.

Our analysis of expression data during embryogenesis of normal ZWf females and temperature sex reversed ZZf females revealed for the first time differences in gene-driven and temperature-driven female development in a single species. Early in development, prior to gonad differentiation, the initiation of the sex reversal trajectory differs from the genetic female pathway both in the timing and genes involved. As development proceeds, differences in expression patterns become less until the pathways converge on a conserved developmental outcome (ovaries). Our ability to compare two female types in *P. vitticeps* allowed us to avoid previously intractable confounding factors such as sex or species-specific differences, which provided unprecedented insight into parallel female pathways. We have provided new insight to the conserved evolutionary origins of the labile networks governing environmentally sensitive sex determination pathways. We have identified a suite of candidate genes, which now provide the necessary foundation for functional experiments in the future. This could include pharmacological manipulation of calcium signalling through alteration of intracellular calcium flux and concentration, such as interference with the TRPV4 channel, or via use of calcium chelators and ionophores, in an organ culture system (17,80). Ongoing development of resources for *P. vitticeps* as an emerging model organism may also allow for RNA interference or gene editing experiments, whereby knock-down or knock-out of candidate genes like *CIRBP, JARID2*, and *KDM6B*, could be used to demonstrate their roles in sex reversal (15).

The maintenance of ovarian differentiation seems to require the operation of different pathways in gene and temperature driven female development. This may involve a pathway centrally mediated by *STAT4* in sex reversed *P. vitticeps*, which has not been previously described, so requires additional confirmation with functional experiments. It will be interesting to determine if a role for these genes occurs in other species. Another STAT family gene, *STAT3*, has recently been demonstrated to play a critical role in the phosphorylation of *KDM6B* and subsequent demethylation of the *DMRT1* promoter required for male development in *T. scripta* (17). The involvement of different genes in the same family is intriguing in its implications; while different genes may be co-opted, natural selection may favour gene families with conserved functions even between evolutionarily disparate lineages.

Our data provided insight into the molecular landscape of the cell required to initiate temperature induced sex reversal. This is the first dataset to capture temperature-induced sex reversal in a reptile, and remarkably we have simultaneously implicated all functional candidates that have previously been identified to be involved in TSD across a range of other species (Table 1). Our results also identified novel genes involved with thermosensitive sex determination, and provide corroborative evidence for the CaRe hypothesis (13). Importantly, our work highlights avenues for future studies to conduct functional experiments to definitively identify the genes and pathways implicated here in sex reversal. Observation and manipulation of intracellular calcium concentrations, as has been conducted in *T. scripta* (17), will also be crucial for fully understanding the role of calcium signalling in sex reversal.

**Table 1:**
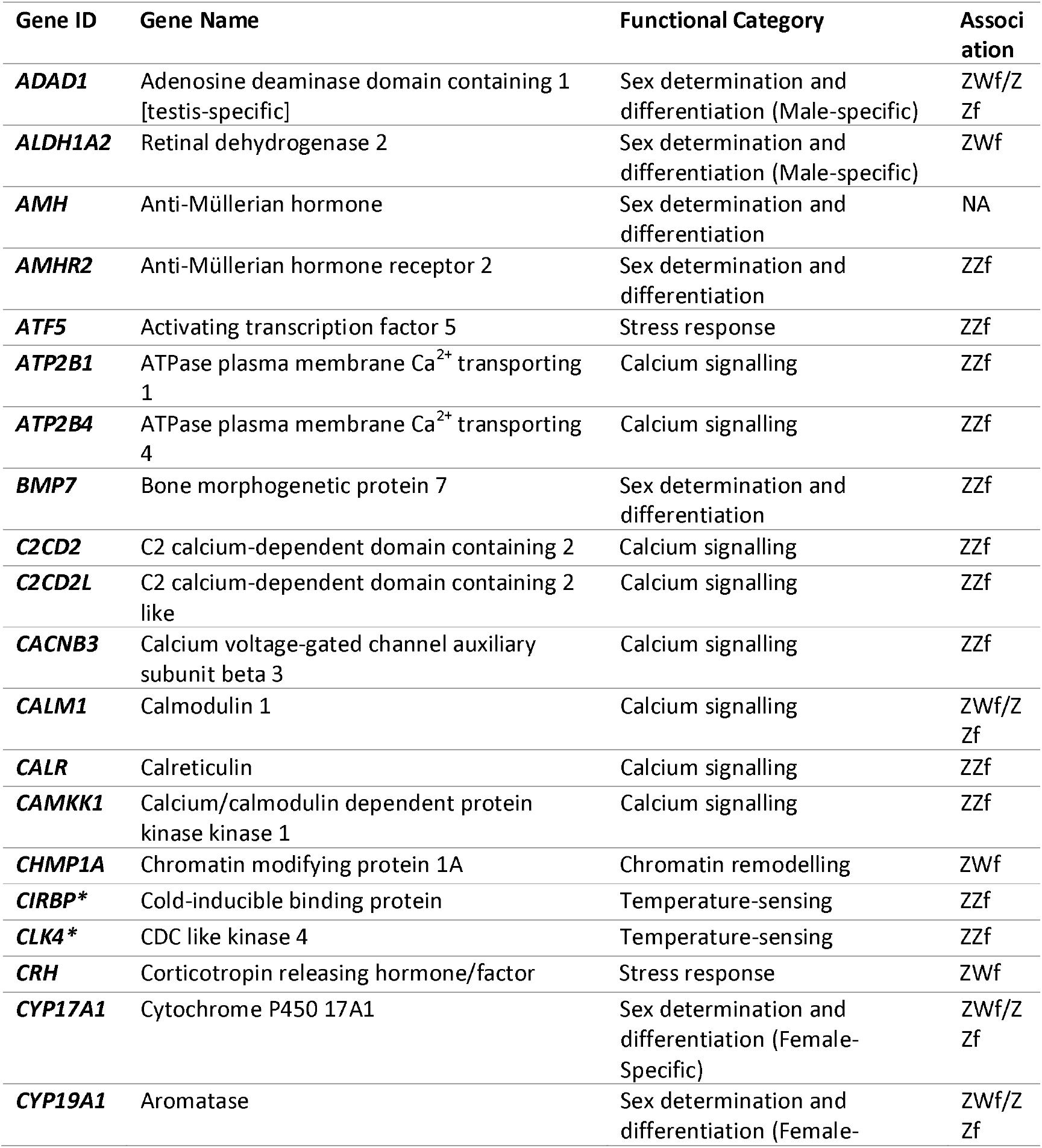
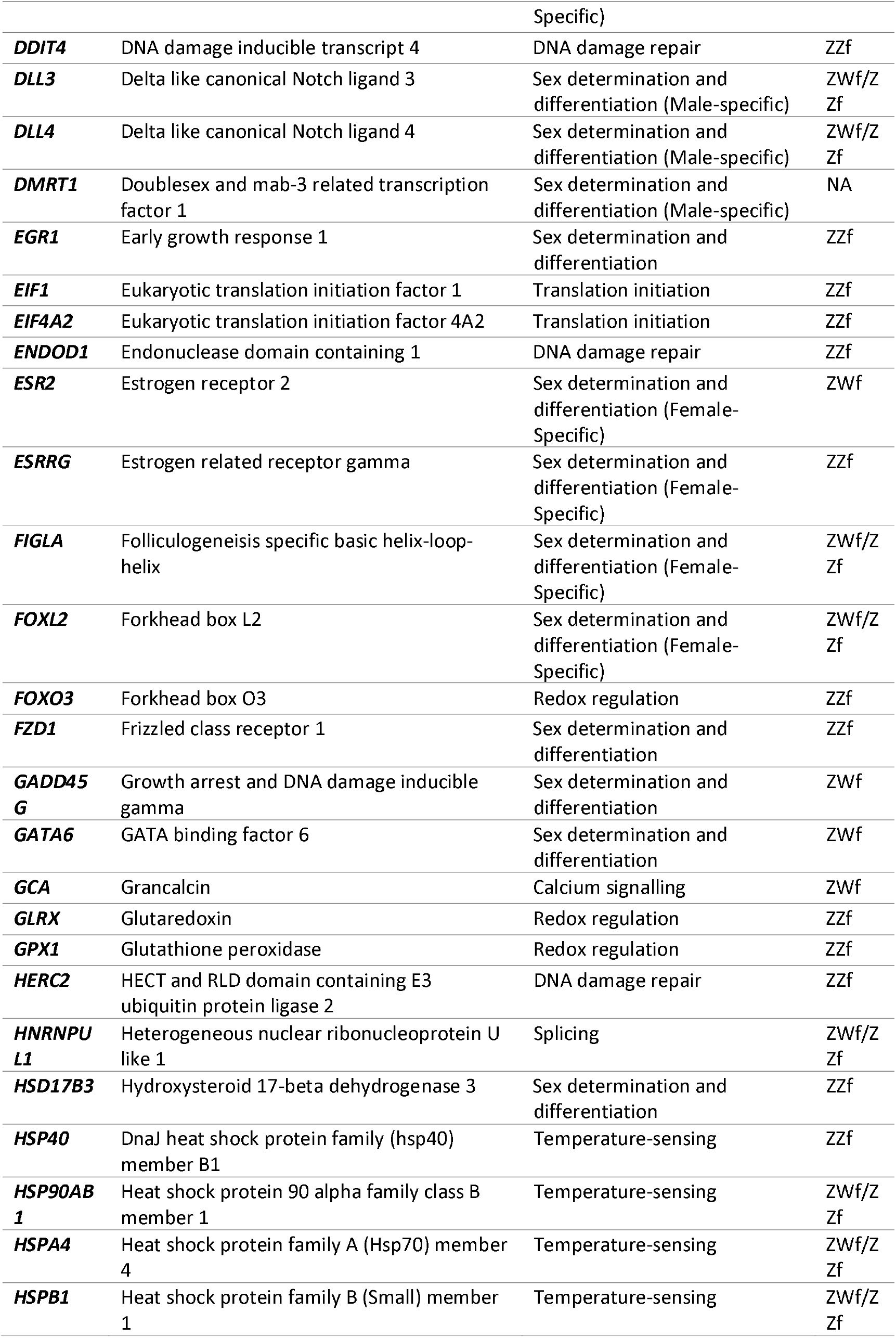
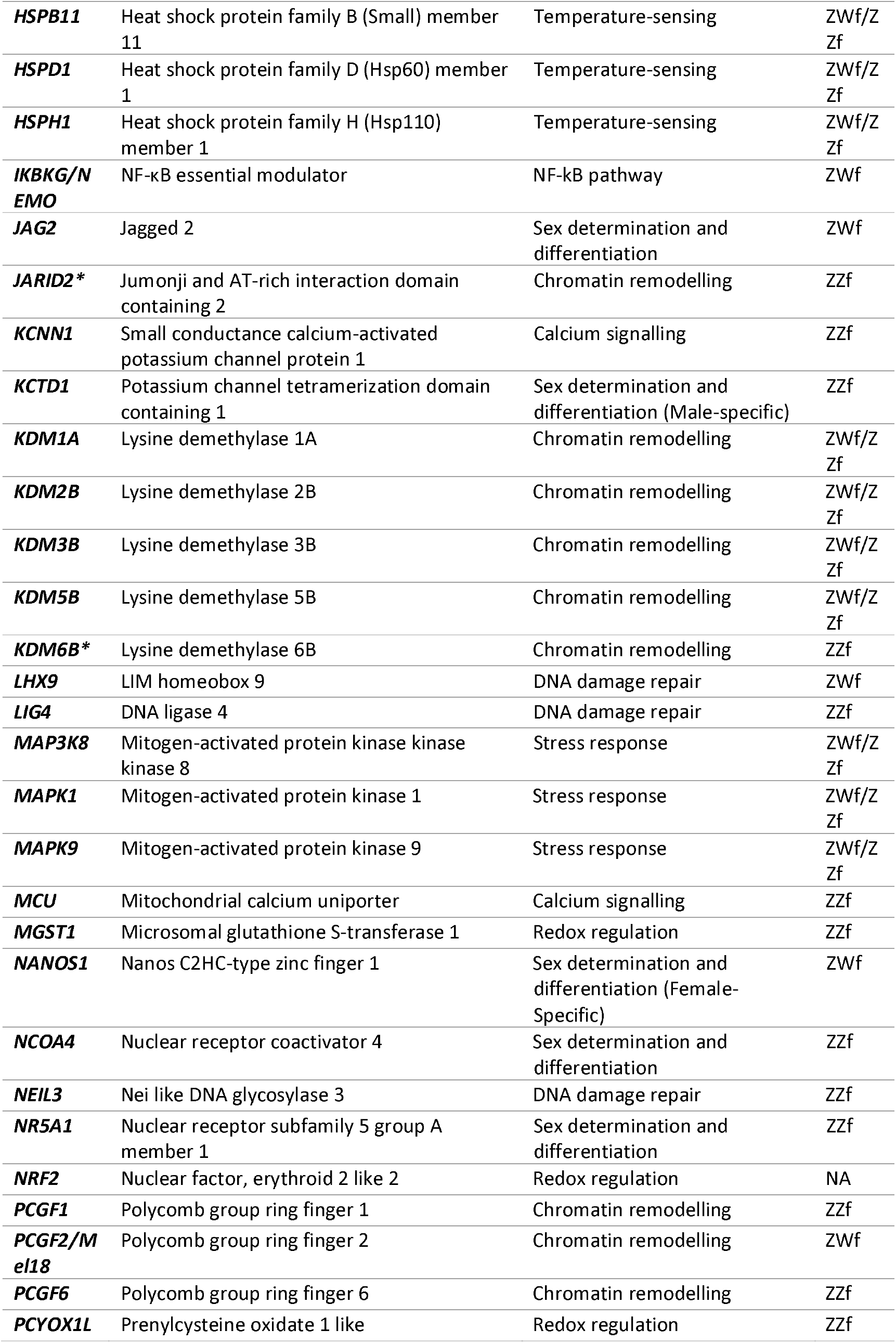
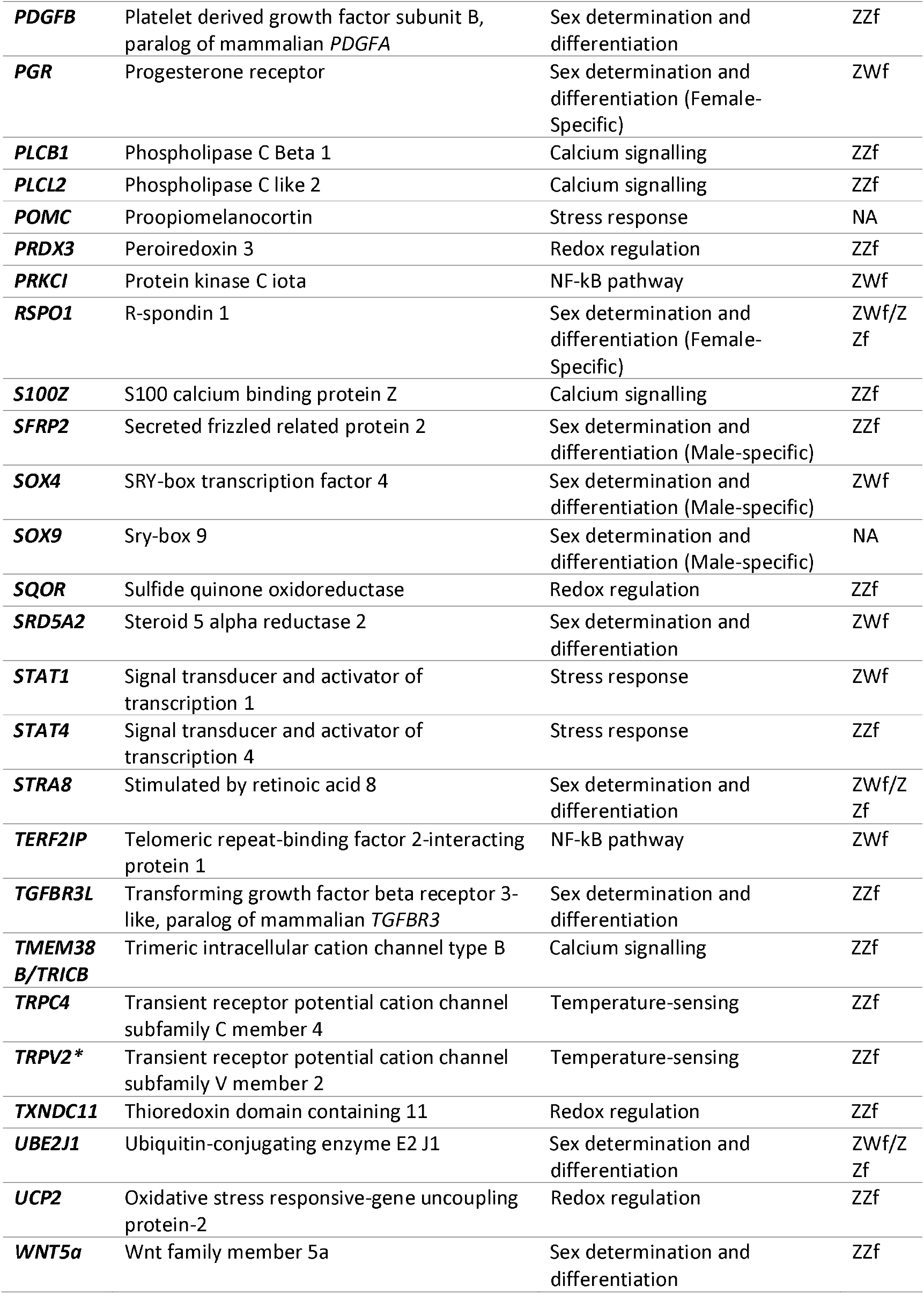
All genes, full gene names, functional categories and associations with either gene (ZWf) or temperature driven (ZZf) female development mentioned in the paper. NA denotes a gene that was mentioned, but was not differentially expressed. Genes with an asterisk are those that have previously been implicated in thermosensitive sex determination cascades, either in *Pogona vitticeps*, or in another reptile species.

Our results highlight the complexity of initiating thermolabile systems. Indeed, it has been suggested that thermolabile sex determination involves system-wide displacement of gene regulation with multiple genes and gene products responding to temperature leading to the production of one sex or the other – a parliamentary system of sex determination (151). We take an intermediate position, arguing for a central role for Calcium-Redox balance as the proximal cellular sensor for temperature, but interacting with other required thermosensitive genes or gene products (e.g. CLK4) to influence ubiquitous signalling pathways and downstream splicing regulation, epigenetic modification and sex gene expression. The level of interaction between each thermosensitive element remains to be explored. For example, if temperature can be sensed by both *TRPV2* and *CLK4*, are both required to initiate sex reversal, or is the signal from only one sufficient? This raises the possibility that no single proximal sensor of the environmental exists, but that several thermosensitive elements early in development must come together to orchestrate alterations in gene expression.

It has been suggested that the products of TRP family genes act as mediators between the temperature signal and a cellular response through Ca^2+^ signalling and subsequent modulation of downstream gene targets (80–82). Notably, different TRP channels are implicated in two alligator species; *TRVP4* in A. mississippiensis, but *TRPV2, TRPC6*, and *TRPM6* in *A. sinsensis*. In *T. scripta, TRPC3* and *TRPV6* are upregulated at male producing temperatures (26°C), while *TRPM4* and *TRPV2* are upregulated at female producing temperatures (31°C), as is the case for *TRPV2* in *P. vitticeps* (79). The diversity of TRP channels recruited for roles in environmental sex determination hints at considerable evolutionary flexibility, perhaps the result of repeated and independent co-option of these channels in TSD species. As may be the case for STAT family genes, the evolution of environmentally sensitive sex determination pathways may involve the use of different genes within gene families that have conserved functions.

Our data also highlights the importance of chromatin remodelling genes in sex reversal in *P. vitticeps. KDM6B* and *JARID2* have been previously implicated sex differentiation in adult *P. vitticeps* (14), embryonic *T. scripta* (15,17) and embryonic *A. mississippiensis* (14). Sex-specific intron retention was observed in TSD alligators and turtle, and was exclusively associated with sex reversal in adult *P. vitticeps* (14). Subsequently, knockdown of *KDM6B* in T. scripta caused male to female sex reversal by removing methylation marks on the promoter of *DMRT1*, a gene critical in the male sex determination pathway (15). *KDM6B* and *JARID2* have also been associated with TSD in another turtle species (*Chrysemys picta*) (126), female to male sex change in the bluehead wrasse, *Thalassoma bifasciatum* (140), and thermal responses in the European bass, *Dicentrarchus labrax* (141).

It is currently unknown if the unique splicing events in *KDM6B* and *JARID2* in adult sex reversed *P. vitticeps* that cause intron retention and presumed gene inactivation, also occur in embryos. Given the high expression of these genes during embryonic development at sex reversing temperatures, it would be surprising if this pattern was observed. We also show a significant role for *CIRBP* as the only other gene, alongside *KDM6B*, to be consistently upregulated during sex reversal in all developmental stages assessed. *CIRBP* is a mRNA chaperone, which could be required to stabilise transcripts of crucial sex specific genes during oxidative, cellular and/or thermal stress. It has been proposed as a novel TSD candidate gene in the turtle, *Chelydra serpentina* (120). This gene remains a promising candidate for mediating thermosensitive responses in TSD more broadly, and its role needs to be explored in more detail.

## Conclusions

The alternative female pathways in *P. vitticeps* demonstrates that there is inherent flexibility in sex determination cascades even within the same species. This is consistent with the idea that, provided a functional gonad is produced, considerable variation in sex determining and differentiation processes at the early stages of development is tolerated under natural selection (151). Perhaps this makes the astonishing variability in sex determination between diverse species less surprising. Our findings provide novel insights, and are a critical foundation for future studies of the mechanisms by which temperature determines sex.

## Materials and Methods

### Animal breeding and egg incubations

Eggs were obtained during the 2017-18 breeding season from the research breeding colony at the University of Canberra. Breeding groups comprised three sex reversed females (ZZf) to one male (ZZ), and three concordant females (ZWf) to two males (Fig. 1). Paternity was confirmed by SNP genotyping (Fig. S2). Females were allowed to lay naturally, and eggs were collected at lay or within two hours of lay. Eggs were inspected for viability as indicated by presence of vasculature in the egg, and viable eggs were incubated in temperature-controlled incubators (±1°C) on damp vermiculite (4 parts water to 5 parts vermiculate by weight). Clutches from sex reversed females (that is, ZZf x ZZm crosses) comprised eggs with only ZZ genotypes. These were initially incubated at 28°C (male producing temperature, MPT) to entrain and synchronise development. After 10 d of incubation, half of the eggs selected at random from each clutch was shifted to 36°C (female producing temperature, FPT). Clutches from ZWf x ZZm crosses were incubated at 28°C throughout the incubation period (Fig. 1A). Sample sizes are given in Fig. 1 and Additional file S7.

### Embryo sampling and genotyping

Eggs from both temperatures were sampled at times corresponding to three developmental stages (6, 12 and 15) (20), taking into account the differing developmental rates between 28°C and 36°C. These stages equate to the bipotential gonad, recently differentiated gonad, and differentiated gonad respectively (19). Embryos were euthanized by intracranial injection of 0.1 ml sodium pentobarbitone (60mg/ml in isotonic saline). Individual gonads were dissected from the mesonephros under a dissection microscope and snap frozen in liquid nitrogen. Isolation of the gonad from the surrounding mesonephros was considered essential for studying transcriptional profiles within the gonad. Embryos from three different ZZf x ZZm clutches from each treatment class (temperature x stage) were selected for sequencing, and randomized across sequence runs to avoid batch effects. Embryos from concordant ZWf x ZZm crosses potentially yield both ZW and ZZ eggs, so these were genotyped using previously established protocols (8,20). Briefly, this involved obtaining a blood sample from the vasculature on the inside of the eggshell on a FTA Elute micro card (Whatman). DNA was extracted from the card following the manufacturer protocols, and PCR was used to amplify a W specific region (8) so allowing the identification of ZW and ZZ samples.

### RNA extraction and sequencing

RNA from isolated gonad samples was extracted in randomized batches using the Qiagen RNeasy Micro Kit (Cat. No. 74004) according to the manufacturer protocols. RNA was eluted in 14 µl of RNAase free water and frozen at −80°C prior to sequencing. Sequencing libraries were prepared in randomized batches using 50 ng RNA input and the Roche NimbleGen KAPA Stranded mRNA-Seq Kit (Cat. No. KK8420). Nine randomly selected samples were sequenced per lane using the Illumina HiSeq 2500 system, and 25 million read-pairs per sample were obtained on average. Read lengths of 2 x 150 bp were used. All samples were sequenced at the Kinghorn Centre for Clinical Genomics (Garvan Institute of Medical Research, Sydney). All sample RNA and library DNA was quantified using a Qubit Instrument (ThermoFisher Scientific, Scoresby, Australia), with fragment size and quality assessed using a Bioanalyzer (Agilent Technologies, Mulgrave, Australia).

### Gene expression profiling

Paired-end RNA-seq libraries (.fastq format) were trimmed using trim_galore with default parameters (v0.4.1; https://www.bioinformatics.babraham.ac.uk/projects/trim_galore/, last access 21-Apr-2020). Trimmed reads were aligned to the *Pogona vitticeps* NCBI reference genome (pvi1.1, GenBank GCA_900067755.1; (142)) using STAR (v2.5.3; (143)), with splice-aware alignment guided by the accompanying NCBI gene (*Pvi1*.*1*) annotation (.gtf format). Likely PCR duplicates and non-unique alignments were removed using samtools (v1.5; (144)). Gene expression counts and normalised expression values (reported in TPM) were determined using RSEM (rsem-calculate-expression; v1.3.1; (145)).

### Identification of non-sex reversed specimens

Normalised transcripts per million (TPM) for a panel of sex-specific genes (*SOX9, AMH, DMRT1, FOXL2, CYP19A1, CYP17A1*) were inspected across the three stages to identify if any samples showed aberrant expression patterns. This approach was also used to determine if any of the stage 12 and 15 samples from the 36°C treatment had not undergone sex reversal by comparing expression levels between ZWf and ZZf embryos; the rate of sex reversal is 96% at 36°C (8) (Fig. S3, Additional file S8). The five samples from clutch 9 exhibited significantly higher expression values for *SOX9, AMH*, and *DMRT1* and represented clear outliers. This was also supported by multidimensional scaling (MDS) plots, so the decision was made to regard the five samples from clutch 9 as aberrant and exclude them from subsequent analyses (Figs S3, S4, Additional file S9). Any ZZf samples with male-like gene expression patterns (high expression for male-specific genes, and low expression for female-specific genes) were considered to have not been reversed (sex reversal is not 100% at 36°C) and were removed (two stage 15 samples).

### Differential expression analysis

Differential expression analysis of ZZf and ZWf transcripts was conducted on raw counts using the EdgeR package (Bioconductor v 3.9 (146)) in R (v 1.2.1335, (147)), following standard procedures outlined in the EdgeR users guide (146,148). Lowly expressed genes, which was applied to genes with fewer than ten counts across three samples, were removed from the raw counts (19,285 genes) so that the total number of genes retained was 17,075. Following conversion to a DGElist object in EdgeR, raw counts were normalised using the upper-quartile method (calcNormFactors function) (149). Estimates for common negative binomial dispersion parameters were generated (estimateGLMCommonDisp function) (148), followed by generation of empirical Bayes dispersion estimates for each gene (estimateGLMTagwiseDisp function) (148,150). A quasi-likelihood binomial generalised log-linear model was fitted (glmQLFit function) and the glmQFTest function was used to compare contrasts within the design matrix (151–155). A P-value cut-off of 0.01 and a log_2_-fold change threshold of 1 or −1 was applied to all contrasts (topTags function) (151). Contrasts were used to assess differential expression between ZZf and ZWf samples across each developmental stage. Raw count (Additional file S10) and expression files (Additional file S11) from this analysis are supplied.

Gene ontology (GO) analysis was conducted for each set of differentially expressed genes using GOrilla (156,157). The filtered count data file (17,075 genes) was used for the background gene set at a P-value threshold of 10^−3^.

### K-means clustering analysis

K-means clustering analysis was performed on normalised counts per million extracted from the DGElist object produced by the initial process of the DGE analysis using edgeR (see above). Counts for each gene were averaged for each treatment group, and the number of clusters was selected using the sum of squared error approach, which was further validated by checking that each cluster centroid was poorly correlated with all other cluster centroids (maximum correlation 0.703 in ZWf clusters, and 0.65 in ZZf clusters). A total of 6 clusters was chosen, and clustering analysis was conducted using the kmeans function in R package stats v3.6.2. Resultant gene lists were sorted by unique and shared genes between clusters with similar trends between ZWf and ZZf (cluster 1 in ZWf, cluster 3 in ZZf, and cluster 3 in ZWf and cluster 5 in ZZf). Both unique and shared genes from each cluster and pairs of clusters (cluster 1 and 3, and clusters 3 and 5) were then analysed for gene ontology (GO) enrichment using GOrilla (156,157). The filtered count data file (17,075 genes) was used for the background gene set at a P-value threshold of 10^−3^.

## Declarations

### Ethics Approval

All procedures were conducted in accordance with approved animal ethics protocols from the University of Canberra Animal Ethics Committee (AEC 17-17).

### Consent for publication

Not applicable

### Availability of data and materials

The raw input files (counts and transcripts per million) that were analysed for this study are available as supplementary files. Raw sequencing data is available under NCBI BioProject PRJNA699086 (Biosample accession IDs SAMN17765903 to SAMN17765941).

### Author Contributions

AG and CEH led and designed the experiment, AG and JMG built the “Sex in Dragons” program of which this is a part. SLW carried out all experimental procedures and differential gene expression analysis. JB created all RNA libraries and carried out all sequencing. IWD generated all data files from sequencing outputs. SW assisted with data analysis, particularly K-means clustering. SLW lead the preparation of the manuscript, with AG, CEH, and JMG. All authors provided feedback on the manuscript and approved the final draft.

## Acknowledgements

We thank Dr Wendy Ruscoe and Jacqui Richardson at the University of Canberra Animal House Facility for their animal husbandry expertise.

## Supplementary Materials

**Additional file S1:** Differentially expressed genes between developmental stages 6 and 12, and 12 and 15 for ZZf and ZWf females generated from EdgeR’s “topTags” function. Results are sorted by log-fold change, with a cut-off of 1 or −1 applied, and a P-value threshold of 0.01. For the stage 6 and 12 comparison, genes with positive log-fold changes are upregulated at stage 12, while for the stage 12 and 15 comparison, genes with a positive log-fold change are upregulated at stage 15. No genes were differentially expressed between stages 12 and 15 in ZZf females.

**Additional file S2:** Gene ontology (GO) enrichment for process and function for the stage 6 and 12 comparison for ZZf and ZWf (Additional file S1). GO enrichment was not possible for the stage 12 and 15 comparison because differentially expressed genes were lacking. GO enrichment was generated from GOrilla (156,157) at a significance threshold of *P* ≤ 0.05.

**Additional file S3:** Differentially expressed genes between ZZf and ZWf for each developmental stage generated from EdgeR’s “topTags” function. Results are sorted by log-fold change, with a cut-off of 1 or −1 applied, and a P-value threshold of 0.01 Genes with positive log-fold changes are upregulated in ZZf embryos, genes with negative log-fold changes are upregulated in ZWf.

**Additional file S4:** Gene ontology (GO) enrichment for process and function generated from GOrilla (156,157) at a significance threshold of *P* ≤ 0.05 for differentially expressed genes at stages 6 and 12 for ZZf and ZWf samples (Additional file S3).

**Additional file S5:** Gene list outputs for K-means clustering analysis (n = 6), and comparative information for matched clusters between ZZf and ZWf (ZZC1 and ZWC1, and ZZC2 and ZWC4) including genes that are unique and shared between each cluster.

**Additional file S6:** Gene ontology (GO) process and function enrichment for genes in ZZf C1, and genes shared between ZZC5 and ZWC3 generated from GOrilla (156,157) at a significance threshold of *P* ≤ 0.05.

**Additional file S7:** Summary of all embryonic gonad samples sequenced for this study, including incubation temperature, genotype, parental cross, developmental stage, and clutch for each sample. Unique sample identifiers are matched to those used in raw data inputs (Additional file S10, S11).

**Additional file S8:** Outputs from pairwise T-tests conducted between stage 15 (n = 7) and stage 15 ZZf samples suspected not to have undergone sex reversal (n = 2). The normalised transcripts per million (TPM) for six genes, three male (*AMH, DMRT1, SOX9*) and three female genes (*FOXL2, CYP19A1, CYP17A1*) were used. The two samples suspected of not undergoing sex reversal show significantly different expression levels for four of these genes, and differences just above the significance threshold of ≤0.05 for the other two genes. On this basis, these two samples were removed from further analysis.

**Additional file S9:** Outputs from one way analysis of variance (ANOVAs) between all clutches (clutch 1, 2, 3, 6, and 9) across each developmental stage (6, 12 and 15) for a panel of sex specific genes (*AMH, SOX9* and *DMRT1*) to determine whether clutch 9 exhibits aberrant expression levels. The normalised transcripts per million (TPM) generates from the EdgeR pipeline described in the materials and methods was used. Based on the results from this analysis, five samples from clutch 9 were excluded from further analysis.

**Additional file S10:** Raw counts for all samples (n = 39) for all genes (n = 19,284) prior to any filtering or sample removal. Sample ID labels correspond to incubation temperature (36 or 28), maternal genotype/maternal homozygosity/maternal heterozygosity (ZZf or ZWf), sample stage (s1 = stage 6, s2 = stage 12, s3 = stage 15), clutch number (c1, c2, etc.), and replicate ID (e.g., “a” denotes the sample was the first replicate for that sampling point). Sample data is also available in Additional file S7.

**Additional file S11:** Raw expression values (TPM, transcripts per million) all samples (n = 39) for all genes (n = 19,284) prior to any filtering or sample removal. Sample ID labels correspond to incubation temperature (36 or 28), maternal genotype/maternal homozygosity/maternal heterozygosity (ZZf or ZWf), sample stage (s1 = stage 6, s2 = stage 12, s3 = stage 15), clutch number (c1, c2, etc.), and replicate ID (e.g., “a” denotes the sample was the first replicate for that sampling point). Sample data is also available in Additional file S7.

**Fig S1:**
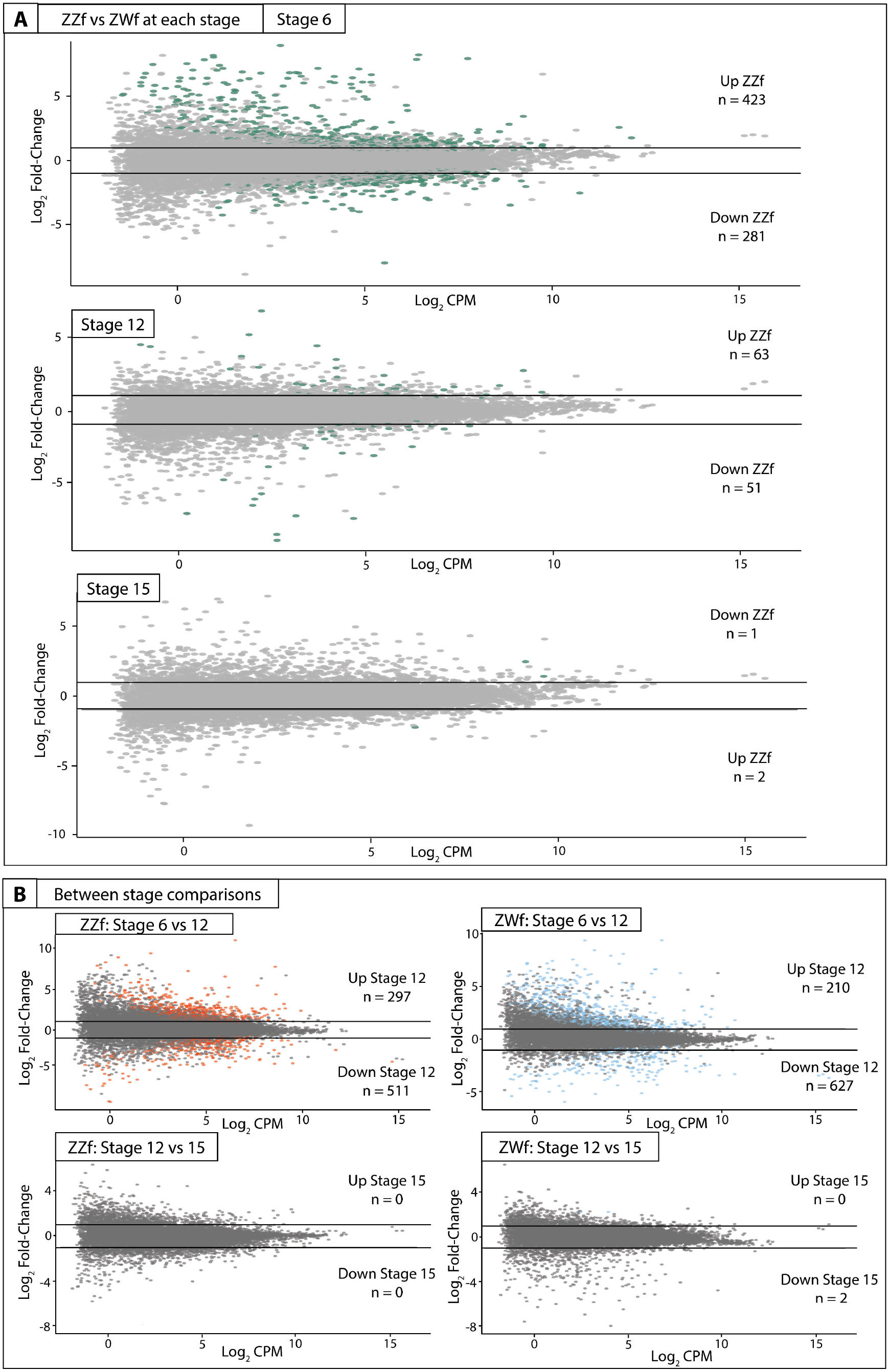
MA plots of read counts per gene from differential expression analysis conducted between ZZf and ZWf (**A**) and comparisons between stages for both ZZf and ZW (**B**). Differentially expressed genes (*P* values ≤ 0.01, log_2_-fold change of 1, −1) are coloured (colour indicative of significant fold change), and the total number of genes are indicated in each plot. Grey indicates no differential expression, horizontal lines indicate log_2_-fold changes of 1, −1. CPM: normalised counts per million

**Fig S2:**
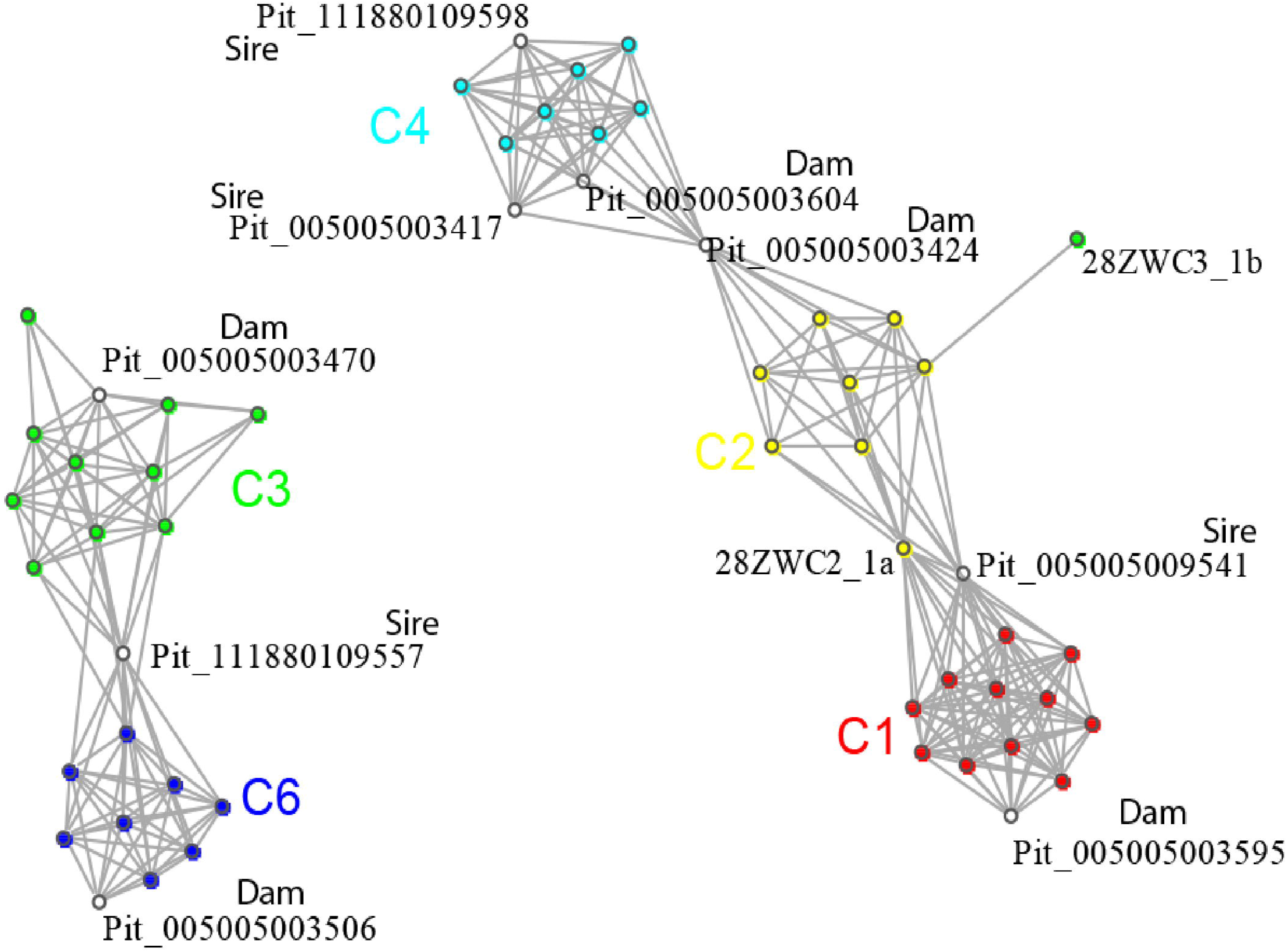
Network analysis of parental and offspring SNPs to confirm paternity of clutches used in this experiment. SNP data was generated by Dart sequencing, a reduced genome representation sequencing method at Diversity Arrays Technology, University of Canberra.

**Fig S3:**
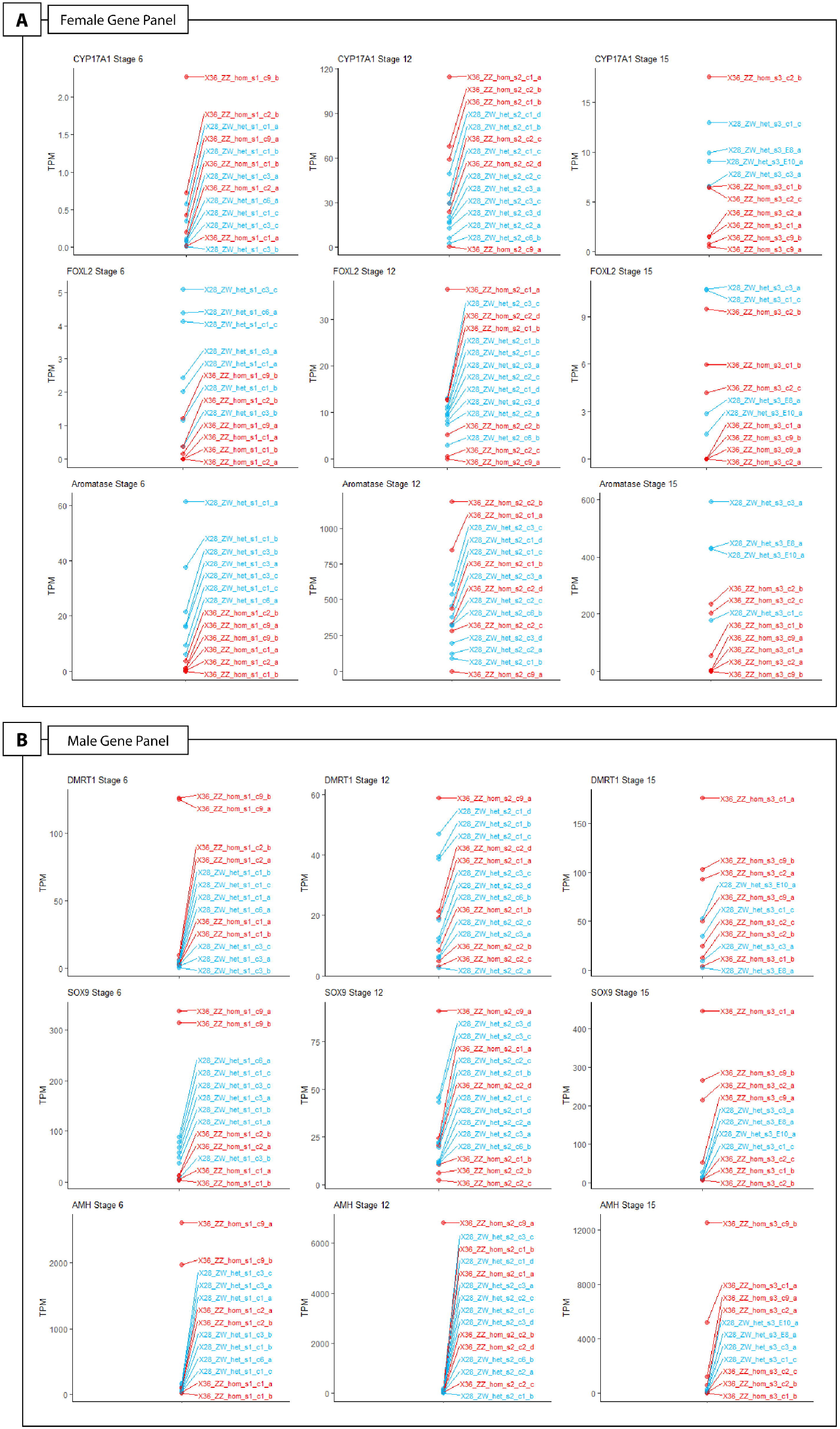
Expression (TPM, transcripts per million) of female-specific genes (*CYP17A1, FOXL2, CYP19A1;* panel A) and male-specific genes (*DMRT1, SOX9, AMH*; panel B) across three developmental stages (6, 12, 15) (19,20) for all samples to aid in the identification of samples with aberrant expression patterns. Samples from later developmental stages that exhibit low expression of female-specific genes are likely to have not undergone sex reversal. Sample ID labels correspond to incubation temperature (36°C or 28°C in red or blue respectively), maternal genotype/maternal homozygosity/maternal heterozygosity (ZZf or ZWf), sample stage (s1 = stage 6, s2 = stage 12, s3 = stage 15), clutch number (c1, c2, etc.), and replicate ID (e.g., “a” denotes the sample was the first replicate for that sampling point).

**Fig S4:**
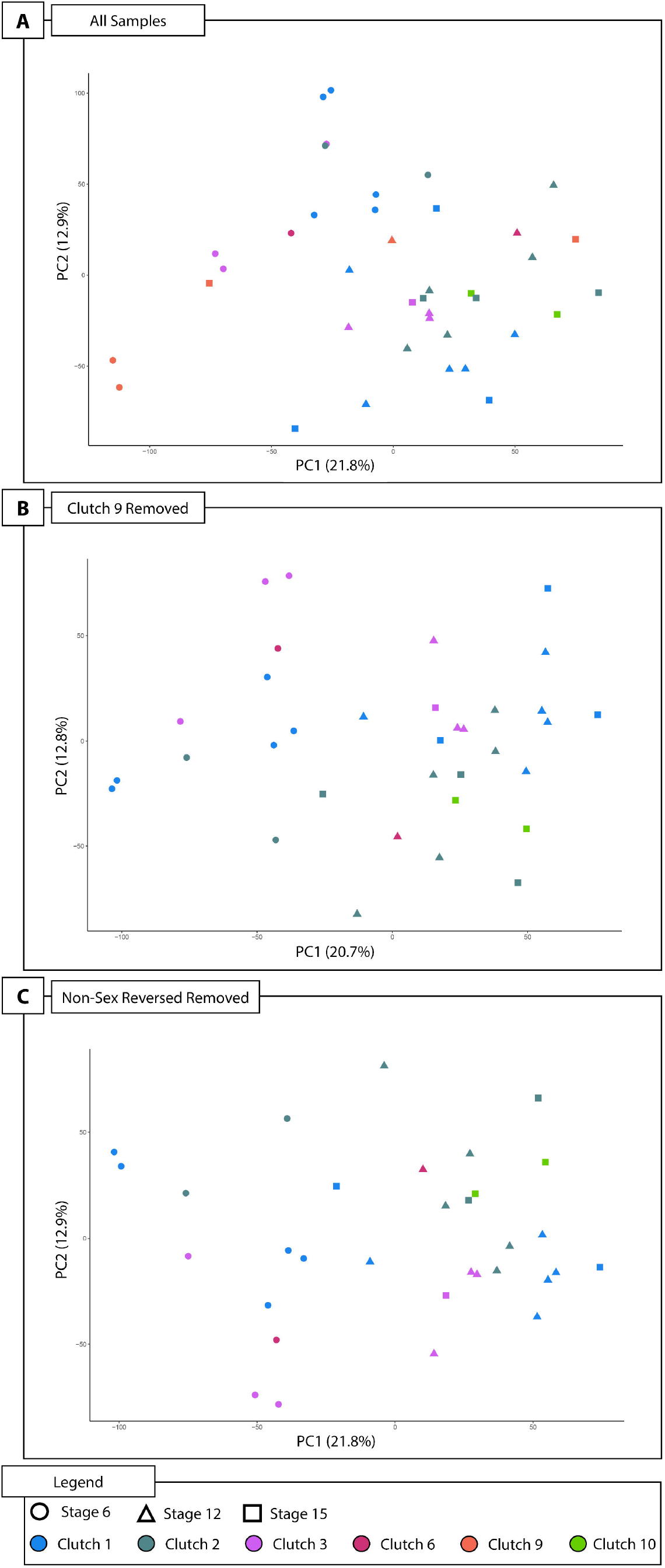
Principal components analysis (PCA) plots performed on normalised counts per million for filtered genes following the EdgeR pipeline described in the materials and methods section. (*A*) PCA of all samples (n = 39) (*B*) PCA of samples with clutch 9 removed (n = 32) (*C*) PCA of samples with clutch 9 samples removed and two samples that had not undergone sex reversal. This is the final dataset upon which all analysis was performed (n = 30).

